# Structural basis for synergistic antibody protection against the essential malaria invasion complex protein RIPR

**DOI:** 10.64898/2025.12.16.693247

**Authors:** Barnabas G. Williams, Jordan R. Barrett, Josefin Bartholdson Scott, Cassandra A. Rigby, Matteo Cagiada, Doris Quinkert, Kirsty McHugh, Anna Huhn, Sean A. Burnap, Camille Gourjault, Francesca Byrne, Sai Sundar Rajan Raghavan, Ana Rodrigues, Laura Bergamaschi, Beatrice Balzarotti, Simon Watson, Noah Miller, Lloyd D. W. King, Francesca R. Donnellan, Camilla A. Gladstone, Jemima Paterson, Stefania Scalabrino, Sarah E. Silk, Jo Salkeld, Angela M. Minassian, Katherine Skinner, Weston B. Struwe, Charlotte M. Deane, Stephen T. Reece, Andrew B. Ward, Simon J. Draper

## Abstract

*Plasmodium falciparum* RH5-interacting protein (RIPR) is central to the essential PTRAMP-CSS-RIPR-CyRPA-RH5 (PCRCR)-complex; a leading target of blood-stage malaria vaccines. However, mechanisms whereby anti-RIPR antibodies inhibit parasite invasion are poorly understood. Here, we characterise 83 human IgG mAbs from RIPR-vaccinated Kymouse platform mice. Single mAbs have minimal neutralising activity, however, high-level synergistic inhibition is observed with pools of mAbs targeting the RIPR-Tail region. Structural characterisation and molecular dynamics simulations of RIPR-Tail show that mAbs targeting EGF-like domains 6-8 (RIPR^EGF^ ^(6-8)^), but not EGF-like domains 9-10 or the C-terminal domain (RIPR^EGF^ ^(9-10)-CTD^), synergise to constrain the RIPR-Tail conformation. The same antibodies dissociate PTRAMP-CSS from RIPR, thereby enabling anti-RIPR^EGF (9-10)-CTD^ mAbs or anti-CSS sdAbs to bind and potentiate anti-RIPR^EGF (6-8)^ IgG. Addition of these mAbs to IgG from humans immunised with the R78C (RIPR^EGF (7-8)^-CyRPA) candidate vaccine enhances malaria growth inhibition. These data provide a framework to guide next-generation blood-stage malaria vaccine design.

## INTRODUCTION

Malaria disease, caused by the parasite *Plasmodium falciparum*, continues to be a global health problem. There were an estimated 282 million cases in 2024 and 610,000 deaths, with most deaths in children under 5 years in sub-Saharan Africa.^1^ The WHO currently recommends two malaria vaccines, RTS/S/AS01 and R21/Matrix-M^2^, with early analysis reporting that vaccine introduction was associated with substantial reductions in hospital admission of young children with severe malaria.^3^ However, both vaccines target the same pre-erythrocytic sporozoite-stage of the parasite lifecycle via the circumsporozoite protein (CSP). Accordingly, a malaria vaccine that targets multiple stages of the parasite lifecycle, including the asexual blood-stage, could improve vaccine effectiveness by providing the immune system with multiple distinct targets.^4^

*P. falciparum* merozoites invade erythrocytes through a complex series of interactions^5^, many of which are redundant, however, one exception is the parasite’s reticulocyte**-**binding protein homologue 5 (RH5) which forms an essential, non-redundant interaction with basigin (BSG/CD147).^6,7^ The current leading blood-stage vaccine candidate (which targets the full-length RH5 molecule) is called RH5.1/Matrix-M^8^, this recently achieved a vaccine efficacy of 55% against clinical malaria over 6 months in young children in a Phase 2b trial in Burkina Faso.^9^ Whilst a substantial milestone in the field, this result shows that significant improvements could still be made to blood-stage vaccines.

RH5 forms a hetero-pentameric complex with cysteine-rich protective antigen (CyRPA)^10^, RH5-interacting protein (RIPR)^11^, *Plasmodium* thrombospondin-related apical merozoite protein (PTRAMP)^12^ and cysteine-rich small secreted protein (CSS)^13,14^, referred to as the “PCRCR-complex”. Each of these proteins are essential for merozoite invasion^14,15^ and are the target of neutralising antibodies^16,17^ or single-domain antibodies (sdAb).^14^ Recent efforts in developing next-generation blood-stage vaccine candidates have either sought to improve on RH5.1 through structure-based vaccine design approaches^18-20^, or by targeting additional components of the PCRCR-complex. One candidate called “R78C” has recently entered Phase 1 clinical testing in the UK and Africa, and comprises RIPR epidermal growth factor (EGF)-like domains 7-8 (RIPR^EGF (7-8)^) fused to CyRPA.^17^

A plethora of information is now available on the RH5 epitope landscape to aid vaccine design approaches. These data come from studies which generated hundreds of anti-RH5 monoclonal antibodies (mAbs) from samples taken either from participants in RH5 vaccine clinical trials^21,22^ or from donors naturally-exposed to malaria in endemic regions.^23^ In addition, a number of high-resolution structures of neutralising anti-CyRPA mAbs are also available.^24-26^ In stark contrast, the epitope data for RIPR, PTRAMP, and CSS are sparse, thus greatly limiting rational vaccine design efforts. RIPR is a large multi-domain protein containing 10 EGF-like domains. Six of these EGF-like domains, RIPR^EGF (5-10)^, and an adjoining C-terminal domain (CTD) extend away from the main body of the RIPR protein to form the “RIPR-Tail”.^27^ The R78C vaccine candidate was designed primarily on data derived from serological analysis of polyclonal anti-RIPR IgG from vaccinated animals and a small number of mouse mAbs.^17^ Moreover, despite its relatively large size (∼120 kDa), only a handful of mouse mAbs against RIPR have been reported^17,28^; of these, few show any *in vitro* growth inhibitory activity (GIA) against *P. falciparum* and typically with EC_50_ values close to 1 mg/mL (2-3 orders of magnitude less effective than the most potent anti-RH5 mAbs).^21,29^ Furthermore, existing cryo-EM structures of RIPR are either low resolution^30^ or lack the RIPR-Tail^27^ (containing known neutralising epitopes^17^), and neither study elucidated the structures of neutralising antibodies, thus limiting our understanding of this vaccine target.

Consequently, although the above studies have provided valuable insights, the relationships that underlie antibody recognition of RIPR and functional growth inhibition remain poorly understood. In this study, we therefore leveraged the humanised Kymouse platform^31^ to generate human mAbs against RIPR, and characterised a large panel of clones to define the determinants of antibody functional potency.

## RESULTS

### GIA-negative synergistic anti-RIPR antibodies bind within the RIPR-Tail

We previously reported that purified polyclonal IgG, raised by vaccination of animals with full-length RIPR (FL-RIPR), could show high-level *in vitro* GIA against *P. falciparum* parasites.^17,32^ This GIA could be completely “reversed” in the GIA assay by addition of recombinant RIPR^EGF (5-8)^, thereby quenching the functional IgG and suggesting that neutralising epitopes lie within RIPR^EGF (5-8)^ and that antibodies against this region are required for GIA. In line with these data, the few known mouse mAbs to show any detectable GIA bind within RIPR^EGF (5-8)^.^17,28^ However, the potency of these antibodies (such as mouse mAbs RP.012 and RP.021) is extremely modest in contrast to the polyclonal vaccine-induced anti-FL-RIPR IgG.^17^ Consequently, we hypothesised that antibodies binding regions of RIPR outside of RIPR^EGF (5-8)^ may not show GIA on their own but may act synergistically with anti-RIPR^EGF (5-8)^ IgG to give higher overall GIA. We thus depleted polyclonal anti-FL-RIPR rabbit IgG of anti-RIPR^EGF (5-8)^ IgG and confirmed that this anti-FL-RIPR^ΔEGF (5-8)^ IgG had no detectable GIA (in line with our previous data). Combining this depleted polyclonal IgG with mAb RP.012 (that binds within RIPR^EGF (6-7)^)^17^ increased the potency of RP.012 from 40% to 80% (**Figure 1A**). These data thus demonstrated that GIA-negative synergistic antibodies were indeed present, potentially in the RIPR-Body and/or RIPR^EGF (9-10)-CTD^ region of the RIPR-Tail. To investigate this, we expressed a series of proteins spanning the RIPR-Tail, each truncated from the N-terminal end by a sequential EGF-like domain (**Figures 1B, S1A**). We next used these proteins to screen for

**Figure 1.**
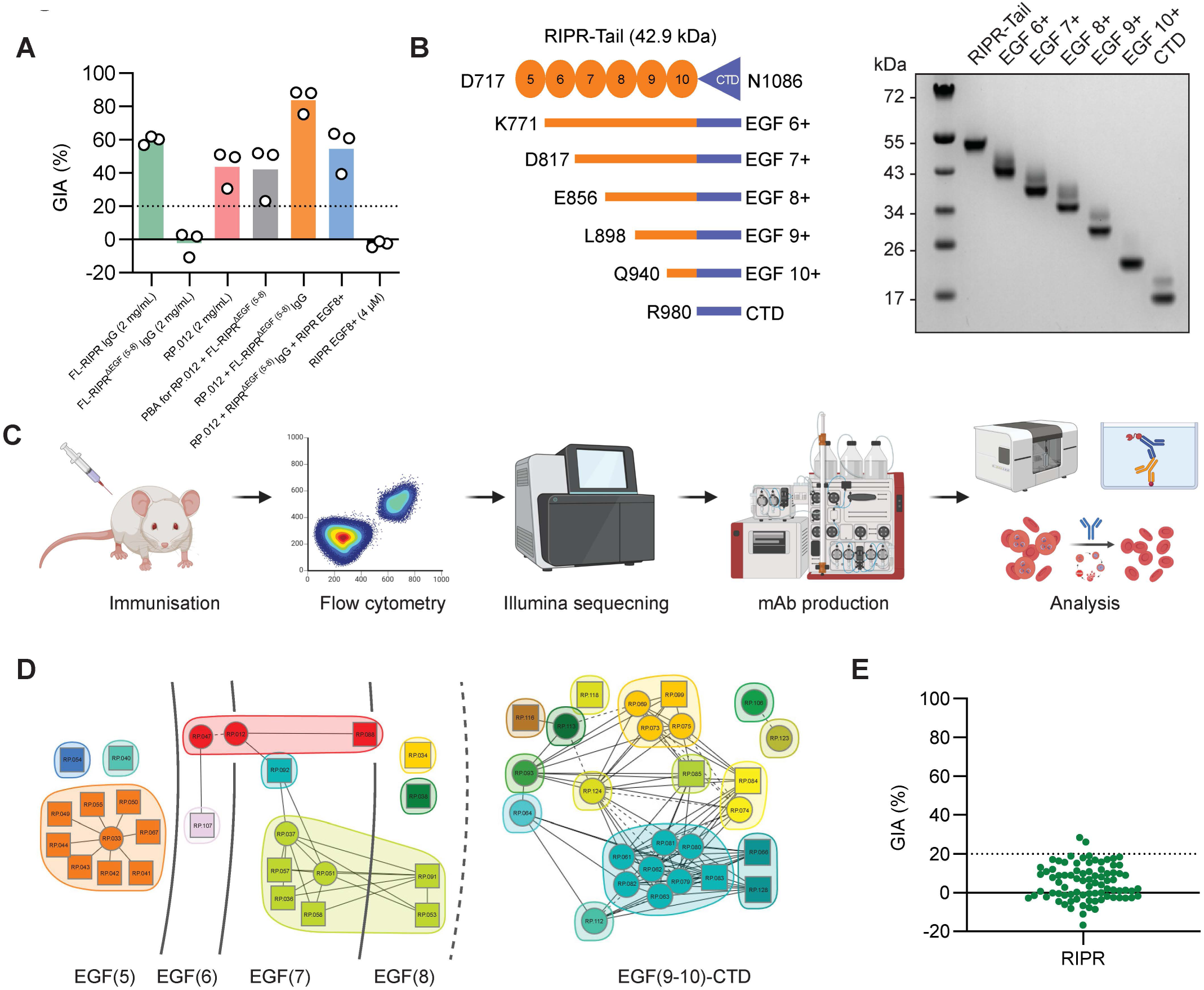
Characterisation of anti-FL-RIPR polyclonal IgG and monoclonal antibodies. **A**) Analysis of anti-RIPR antibodies for *in vitro* GIA against 3D7 clone *P. falciparum* parasites. Samples included: i) Total IgG, purified from sera of rabbits vaccinated with FL-RIPR, at 2 mg/mL concentration either unmodified [FL-RIPR IgG] or depleted of anti-EGF (5-8) IgG [FL-RIPR^ΔEGF (5-8)^ IgG] (green bars); ii) mAb RP.012 at 2 mg/mL (red bar); iii) mAb RP.012 + FL-RIPR^ΔEGF (5-8)^ IgG alone (orange bar) or with addition of RIPR EGF 8+ protein (blue bar). Test group IgG, mAb or protein concentrations are equal to those in control groups. The grey bar shows the GIA predicted by the Bliss’s additivity model (PBA) for the combination of RP.012 mAb and depleted IgG. Black bars show RIPR EGF 8+ protein control at 4 μM final concentration. Dots show N=3 independent repeats and bars the mean. **B**) Diagram and reduced SDS-PAGE gel of the RIPR-Tail and truncation proteins. Orange circles represent EGF-like domains, blue triangle the C-terminal domain (CTD). Truncations of the RIPR-Tail are shown below with the N-terminal amino acid listed on the left and truncation nomenclature on the right. **C**) Schematic showing outline of experimental work for anti-FL-RIPR mAb generation. Preliminary analysis included ELISA, epitope mapping using the Carterra LSA platform, and the GIA assay. **D**) Community network plot illustrating the competitive relationship between 51 vaccine-induced anti-RIPR-Tail mAbs as defined by HT-SPR overlaid with EGF-like domain and CTD binding data derived independently by ELISA (**Data S2**). Individual mAbs are represented as nodes and communities as colours. Solid lines between nodes indicate bidirectional competition. Dashed lines between nodes indicate unidirectional competition. Square nodes indicate mAbs that were excluded as either a ligand or an analyte. Dashed line after EGF (8) indicates the separation between two independent experiments. **E**) GIA assay data, performed as panel (A), of N=83 anti-FL-RIPR IgG mAbs all tested at 1 mg/mL. Each point represents a single mAb. GIA levels below 20% (dotted line) in (A) and (E) are regarded as negative.

GIA-reversal of the synergising activity observed using the FL-RIPR^ΔEGF (5-8)^ IgG, and identified that this synergy could be reversed by addition of 4 μM RIPR EGF 8+ protein, a truncation comprising amino acids E856 – N1086 of RIPR (**Figure 1A,B**). These data thus indicated that the polyclonal anti-FL-RIPR IgG contains GIA-negative antibodies that i) have epitopes within the EGF (8-10)-CTD region of the RIPR-Tail, and that ii) can synergise with GIA-positive anti-RIPR^EGF (5-8)^ IgG.

### Mapping the antibody epitope landscape along the RIPR-Tail

To better understand this phenomenon, we first sought to resolve the epitope landscape across RIPR, with a focus along the RIPR-Tail. We thus immunised Kymouse Platform mice with 10 µg doses of either FL-RIPR protein or the pre-formed recombinant RCR-complex. Intramuscular immunisations were given on days 0, 14, 42, and 56 with spleens harvested on day 61. Anti-RIPR and anti-RCR IgG responses were measured by end-point ELISA (**Figures S1B,C**) and the sera from the top three responding animals were pooled and evaluated by GIA assay, confirming neutralising polyclonal antibodies were present (**Figure S1D**).

Splenocytes from the three highest responding mice were processed and antigen-specific B cells were sorted using either biotinylated FL-RIPR conjugated to streptavidin-PE, or the biotinylated RIPR-Tail components RIPR^EGF (5-8)^ and RIPR^CTD^, conjugated to streptavidin-PE or streptavidin-AF488, respectively. Single cell Ig gene sequencing generated 900 high-quality paired VH+VL sequences, from which 119 unique sequences were chosen aiming to maximise the diversity of antibody clones: 59 targeting FL-RIPR, 29 targeting RIPR^CTD^, and 27 targeting RIPR^EGF (5-8)^. Recombinant mAbs were subsequently cloned as human IgG1, produced at 10 mL scale in Expi293 cells and tested for binding to FL-RIPR by ELISA. 26/119 mAbs showed no binding to FL-RIPR by ELISA and a further 9 could not be expressed to sufficient levels, leaving a final panel of 83 unique anti-FL-RIPR mAbs for further analysis, termed RP.033 to RP.133 (**Figure 1C**).

Kinetic data for each mAb were gathered using the high-throughput surface plasmon resonance (HT-SPR) Carterra-LSA platform (**Data S1; Figure S1E**), all but one mAb had <10 nM affinity. However, epitope binning of the mAb panel on the same platform proved challenging due to some apparent cross reactivity between anti-RIPR^EGF (5-10)^ mAbs, combined with sensitivity of the mAb panel to the low pH regeneration step required by the HT-SPR method. Therefore, we next used the truncated RIPR-Tail protein panel (**Figure 1B**) in an ELISA-based screen to determine where mAb binding is lost, thereby providing a low-resolution location for each mAb epitope on the RIPR-Tail or RIPR-Body (**Data S1,S2**). Sixteen mAbs bound to the RIPR-Body whereas 68 bound to domains in the RIPR-Tail. These data were subsequently paired with two smaller HT-SPR epitope binning experiments designed to mitigate the challenges of EGF-like domain cross-reactivity (**Figures S1F,G**), to generate an epitope map of the RIPR-Tail (**Figure 1D**). The antibody communities identified by HT-SPR mostly aligned to the EGF-like domain binding bins determined by ELISA, however, some cross-domain binding competition was seen as expected due to the size of a mAb relative to a single EGF-like domain. Overall, 11 mAbs bound RIPR^EGF (5)^; 22 bound across RIPR^EGF (6-8)^; and 34 bound RIPR^EGF^ ^(9-10)-CTD^ by ELISA; of these 51 could be binned into communities by HT-SPR.

### Antibodies targeting RIPR-Tail synergise to mediate parasite growth inhibition

To understand functionality of these anti-RIPR antibodies, we next measured *in vitro* GIA against 3D7 clone *P. falciparum* parasites. Each anti-RIPR mAb was first screened in a single point GIA assay at 1 mg/mL. Notably, no mAb exceeded 30% GIA in this assay (**Figure 1E**) with almost all showing <20% (negligible) GIA. We also performed titration curves, starting at even higher concentration, for three mAbs which bind within RIPR^EGF (5-8)^ (RP.034, RP.035, and RP.047), and included the original RP.012 mouse mAb as a control. RP.012 showed modest GIA of ∼40% at 2 mg/mL, but the other three mAbs showed no GIA (**Figure S1H**). We therefore concluded that, despite anti-FL-RIPR polyclonal IgG showing levels of GIA up to ∼80%, that anti-RIPR IgG mAbs individually show modest, if any, neutralising activity.

We thus hypothesised that combinations of anti-RIPR antibodies may be responsible for the GIA observed with polyclonal IgG. In light of our serological data (**Figure 1A**), we focussed on mAbs that bind within RIPR-Tail and designed a pairwise screen at a fixed concentration of antibodies to determine whether inter-domain or intra-domain interactions were important for GIA. Bliss’s additivity^32^ was used to predict the GIA of mAb combinations, although minimal GIA was predicted given every mAb in the panel, except RP.012, showed little or no GIA when tested alone. The results, however, showed that synergistic interactions occurred across the RIPR-Tail (**Figures 2A, S2A,B**). Distinct zones of synergy were observed, in particular when mAbs targeting RIPR^EGF (6-8)^ were combined in a pairwise manner with either i) each other, or ii) with mAbs targeting RIPR^EGF (9-10)-CTD^. In contrast, no synergistic activity was observed for antibodies binding RIPR^EGF (5)^, an EGF-like domain previously associated with neutralising epitopes.^17^ Very little inter-domain antagonism was observed, aside from some occurring with a single anti-RIPR^EGF (6)^ mAb (RP.012). Interestingly, mAbs binding within the RIPR^EGF (9-10)-CTD^ region did not behave synergistically to mediate GIA with each other, although they could improve the GIA of mAbs targeting RIPR^EGF (6-8)^. For example, RP.035, an anti-RIPR^EGF (8)^ mAb, showed minimal GIA when tested alone, but clear synergy with anti-RIPR^EGF (9-10)-CTD^ mAbs such as RP.063 and RP.085 (**Figure 2B**). Taken together with the earlier serological data, these results strongly suggested that RIPR^EGF (6-8)^ forms a “neutralisation zone” of epitopes against which polyclonal IgG responses can be GIA-positive, typically achieved by synergistic interactions. In contrast, RIPR^EGF (9-10)-CTD^ forms a “synergy zone” of epitopes against which polyclonal IgG responses by themselves are GIA-negative but which can, nonetheless, potentiate the GIA mediated by an anti-RIPR^EGF (6-8)^ response.

**Figure 2.**
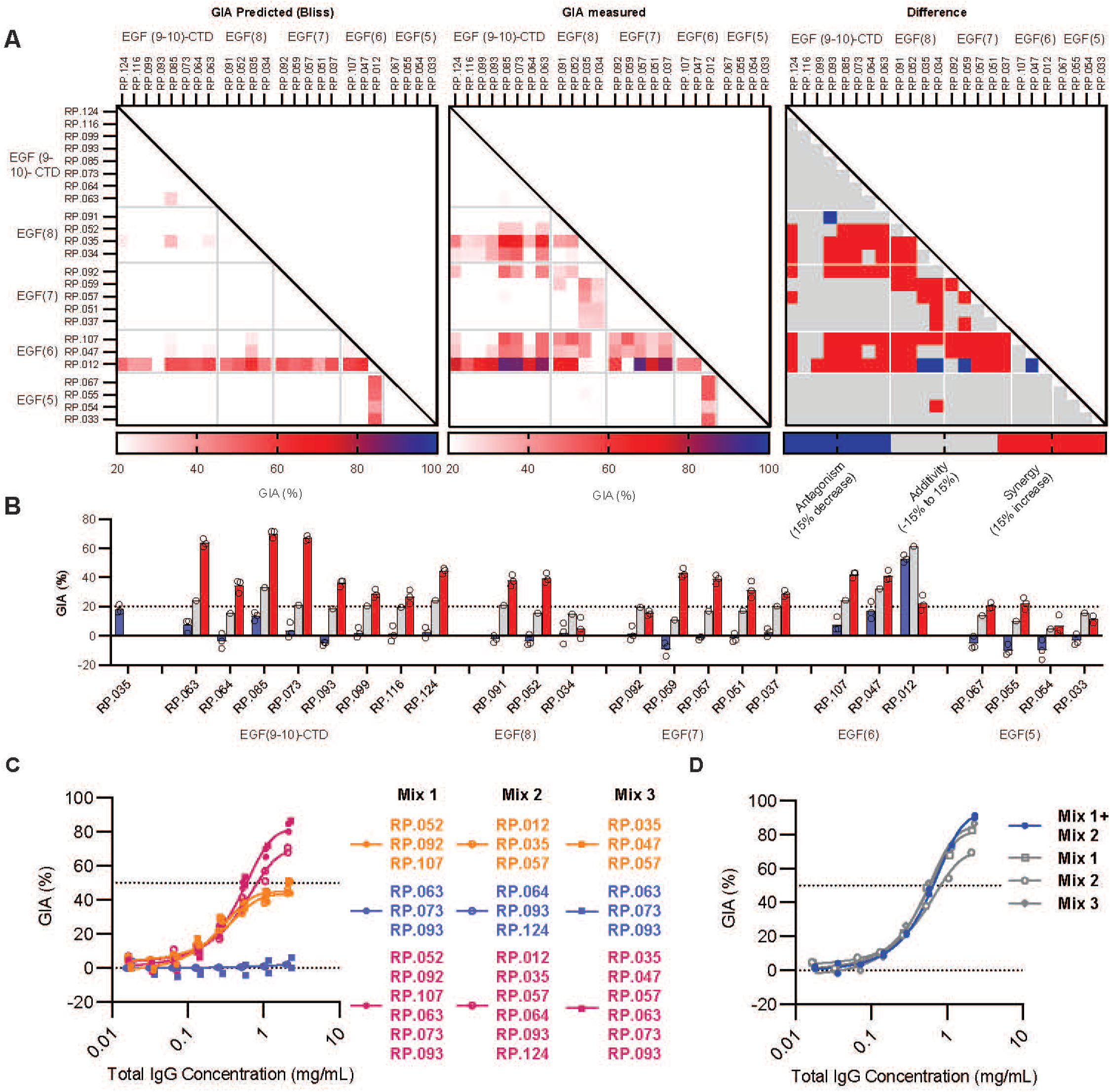
Assessment of intra-RIPR antibody synergy. **A**) Anti-RIPR mAbs were tested for GIA against 3D7 clone *P. falciparum* parasites in pairwise combinations at 1 mg/mL per mAb. Heatmaps show: predicted GIA (%) (assuming Bliss additivity) of each combination (left); measured GIA (%) of each combination (centre); and predicted minus measured GIA (%) (right) where ≥15% GIA difference was classified as synergistic (red) or ≤15% GIA difference as antagonistic (blue), with additivity (within 15% GIA of the predicted value) shown in grey. Data for mAbs are clustered according to their indicated binding domain(s) within the RIPR-Tail. **B**) Example GIA assay dataset for pairwise analysis of anti-RIPR-Tail mAbs with RP.035 (an anti-EGF (8) mAb). Bars show GIA (%) for: second mAb measured alone (blue); the predicted GIA of the second mAb + RP.035 combination assuming Bliss additivity (grey); and the measured result (red). N=3 technical repeats. **C**) GIA assay titrations of anti-RIPR mAb mixtures. Test mAbs were combined in mixtures of 3 or 6 mAbs at an equal ratio (e.g. 1 mg/mL of a triple mAb mixture contains 0.33 mg/mL of each mAb). Orange mAbs target EGF (6-8); blue mAbs target EGF (9-10) and CTD. Red indicates the mix of all six mAbs in the combination. Each data set is fitted with a Richard’s five-parameter dose-response curve with no constraints. Data points show results from N=2 independent repeats. **D**) GIA assay titration performed as per (C) with the 11 anti-RIPR-Tail mAbs combined (in equal ratio) from Mix 1 and Mix 2 shown in blue. GIA data for the hexa-mAb Mixes 1-3 shown in panel (C) are included for clarity in grey.

We next investigated the functional interplay between mAbs targeting the “neutralisation zone” of RIPR^EGF (6-8)^ and the “synergy zone” of RIPR^EGF (9-10)-CTD^. Choosing the most synergistic mAb combinations from the synergy screen, we made three mixes of three non-competing “neutralisation zone” mAbs, and three non-competing “synergy zone” mAbs. RP.093 was included in all “synergy zone” mixes because it was the only mAb identified by ELISA to bind to RIPR^EGF (9)^ in the RIPR^EGF (9-10)-CTD^ group (**Data S2**). The total amount of each mAb in the mix was equal and, with the exception of RP.012, no mAb in these mixes showed GIA when tested in isolation (**Figure S2C**). Here, the mixtures of three mAbs targeting the RIPR^EGF (6-8)^ neutralisation zone produced moderate GIA, plateauing below 50% GIA at 2 mg/mL total concentration, whilst the mixtures of three mAbs targeting the RIPR^EGF (9-10)-CTD^ synergy zone showed no GIA. When all six mAbs were combined, higher GIA was seen, reaching 70-80% at 2 mg/mL (**Figure 2C**) even though each mAb was used at half the concentration of the triple mix comparator; moreover, all antibody mixtures showed similar potency irrespective of mAb composition. We then tested the 11 mAbs from Mix 1 and Mix 2 combined and found that the GIA EC_50_ did not further improve upon that observed for the hexavalent-mAb mixtures (**Figure 2D**). Notably, the six mAb mixtures achieved very similar GIA potency to the original anti-FL-RIPR polyclonal IgG derived from the Kymouse platform mice (**Figure S1D**). These data suggested that functional epitopes span the length of the RIPR-Tail from EGF (6) through to CTD, and that a polyclonal antibody response targeting multiple epitopes underlies the GIA achieved by FL-RIPR immunisation.

### Structure of the RIPR-Tail

The only published high-resolution cryo-EM structure of the RCR-complex does not include the RIPR-Tail (residues D717-N1086)^27^, suggesting inherent flexibility of this string of EGF-like domains or the lack of a large, globular structure for particle alignment in the absence of protein binding partners. Given our data indicated that this uncharacterised region contains the epitopes for functional anti-RIPR antibodies, we sought to structurally characterise this region with synergistic combinations of antibodies using cryo-EM single-particle analysis (cryo-EM SPA). Using a combination of 6 synergistic antibodies targeting EGF domains 6-10 and the CTD (Mix 3 in **Figure 2C**), we were successful in building a model of a complex of RCR bound to RP.047 (RIPR^EGF^ ^(6)^); RP.057 (RIPR^EGF^ ^(7)^); RP.035 (RIPR^EGF^ ^(8)^); and RP.093, RP.073 and RP.063 (RIPR^EGF (9-10)-CTD^) (**Figures 3A, S3A,B; Data S3**). 3D model reconstruction was achieved through combination of two high-resolution cryo-EM maps of RIPR-Tail residues G750-K901 (RIPR^EGF (6-8)^) bound to RP.047, RP.057 and RP.035 at a global resolution of 3.35 Å (PDB ID: 9Q69, EMD-72260) and RIPR-Tail residues V925-N1086 bound to RP.093, RP.073 and RP.063 at a global resolution of 2.97 Å (PDB ID: 9Q6B, EMD-72265). These atomic resolution models, as well as the previously published cryo-EM structure of the RCR-complex containing RIPR residues D34-P716 (PDB ID: 8CDD), were then fitted into a low-resolution 6.1 Å cryo-EM map of the entire complex to generate a full composite model of the RCR:RP.047:RP.057:RP.035:RP.093:RP.073:RP.063 complex (**Figure S3C**). Two small ‘hinge’ regions of the RIPR-Tail comprising half an EGF domain each; residues D717-N749 (RIPR^EGF(5)^) and C902-G924 (RIPR^EGF (9)^) were only visible at low-resolution and unable to be modelled experimentally, likely due to high flexibility. These regions were instead predicted using AlphaFold 3^33^ and aligned with the experimental models and low-resolution map density to build a complete model of the RIPR-Tail.

**Figure 3.**
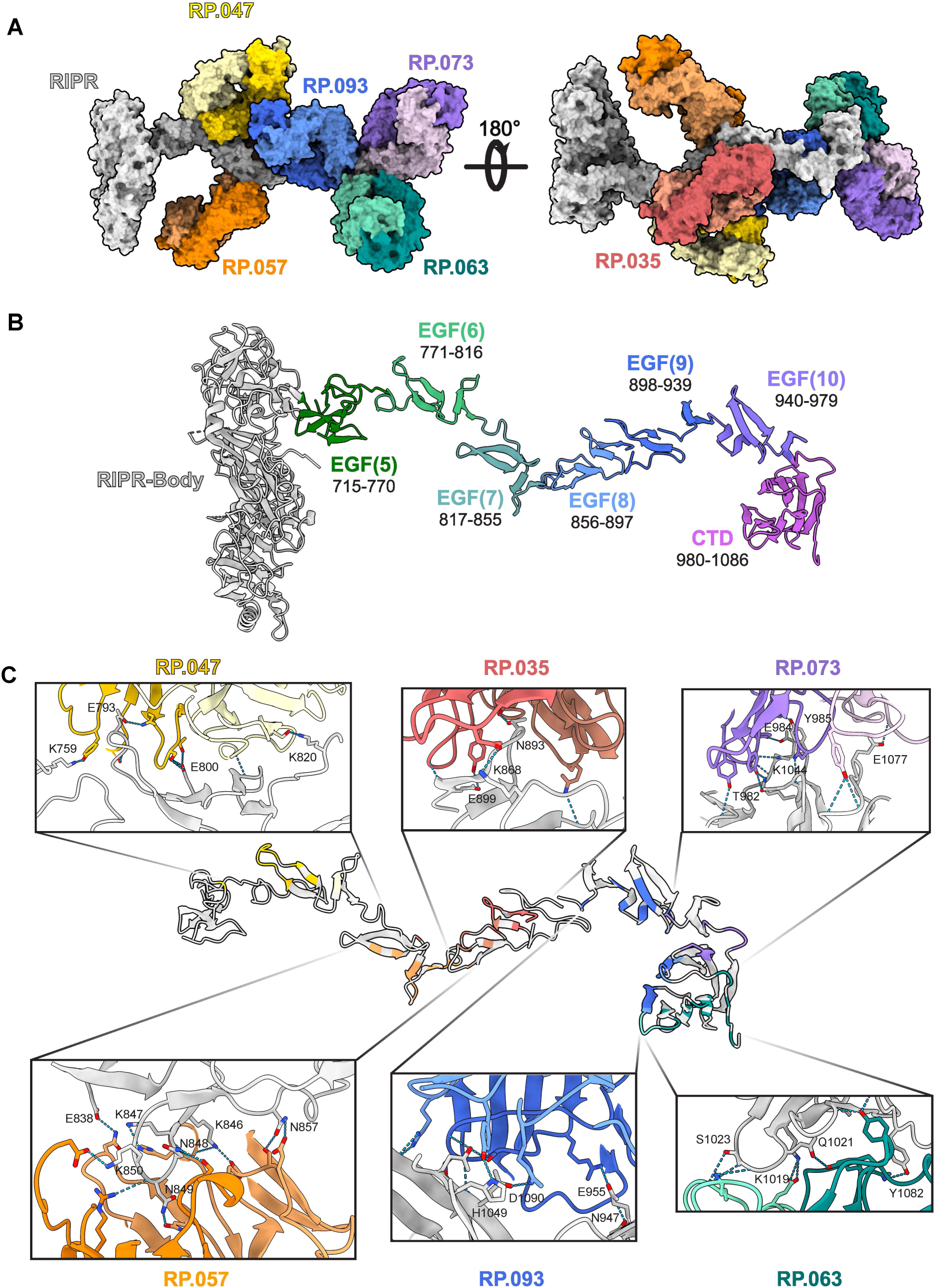
The structure of RIPR-Tail bound to synergistic antibodies. **A**) Structure of the FL-RIPR:RP.047:RP.057:RP.035:RP.093:RP.073:RP.063 complex shown as a surface representation, with the two images flipped 180° relative to one another. RIPR is shown in grey and the six Fabs in colour. **B**) Structure of FL-RIPR residues D34-N1086 shown as a cartoon representation with EGF-like domains 5-10 and CTD highlighted in colour. For clarity, residues D34-C714 which comprise multiple domains are shown as one RIPR-Body domain in grey. **C**) Structure of the RIPR-Tail region (residues V715-N1086) shown as a cartoon representation. The interfacing residues on RIPR with the corresponding Fab (predicted using the interfaces tool with default setting in ChimeraX 10.91) are highlighted in colour. Insets show a close-up of the RIPR:Fab binding interface for each antibody, and RIPR residue side chains predicted to form hydrogen bonding interactions are labelled.

The RIPR-Tail, in the presence of bound antibodies, has an extended, undulating conformation, stretching approximately 160 Å perpendicular to the RIPR-Body (**Figure 3B**). Disulphide-stabilised EGF-like domains comprise most of the RIPR-Tail, connected by short linkers and loops. The RIPR-Tail terminates in the RIPR^CTD^, a globular beta-sandwich domain formed by two 4-strand anti-parallel beta sheets. The Fabs exhibited distinct angles of approach when targeting the RIPR^EGF (6-8)^ domains, predominantly recognising solvent-exposed, unstructured loop regions within specific EGF-like domains (**Figure 3C; Table S1**). RP.035 interfaced entirely with residues in RIPR^EGF (8)^, RP.047 spans RIPR^EGF (5-7)^ with most contacts in RIPR^EGF (6)^, whilst RP.057 predominantly bound RIPR^EGF (7)^ with some contacts within RIPR^EGF (8)^. Of the anti-RIPR^EGF (9-10)-CTD^ antibodies: RP.063 interfaced entirely with residues in RIPR^CTD^; RP.073 bound to RIPR^EGF (10)-CTD^; whereas the RP.093 interface spans RIPR^EGF (9-10)-CTD^ with contacts in all three domains. All three of these antibodies appeared to bind the same side of RIPR^EGF (9-10)-CTD^, leaving one face of the region unbound.

### Anti-RIPR-Tail antibody binding in context of the CSS-RIPR-CyRPA-RH5 complex

Having solved the structure of the RIPR-Tail in complex with a synergistic cocktail of antibodies, we then investigated how these antibodies bind in the context of the wider PCRCR-complex; here RH5 binds to membrane-bound basigin complexes and the RIPR^EGF (9-10)-CTD^ region is anchored to the merozoite membrane via the PC heterodimer.^7,14,34^ We first docked our full-length RCR:RP.047:RP.057:RP.035:RP.093:RP.073:RP.063 composite model to the full-length basigin:MCT1 complex anchored to the erythrocyte membrane (PDB IDs: 4U0Q; 6LZ0). This superimposition of models visualised the binding of antibodies relative to the erythrocyte membrane during a merozoite invasion event (**Figure 4A**). This model of the invasion synapse, in this antibody bound state, predicts that the RIPR-Tail runs parallel to the merozoite and erythrocyte membranes, with the RIPR-Tail presenting an erythrocyte-proximal and merozoite-proximal face ∼150 Å from the erythrocyte membrane. The antibodies binding RIPR^EGF (6-8)^ are orientated parallel to both membranes, whilst the anti-RIPR^EGF (9-10)-CTD^ antibodies bind the erythrocyte-proximal face of the RIPR-Tail, pointing towards the erythrocyte membrane.

**Figure 4.**
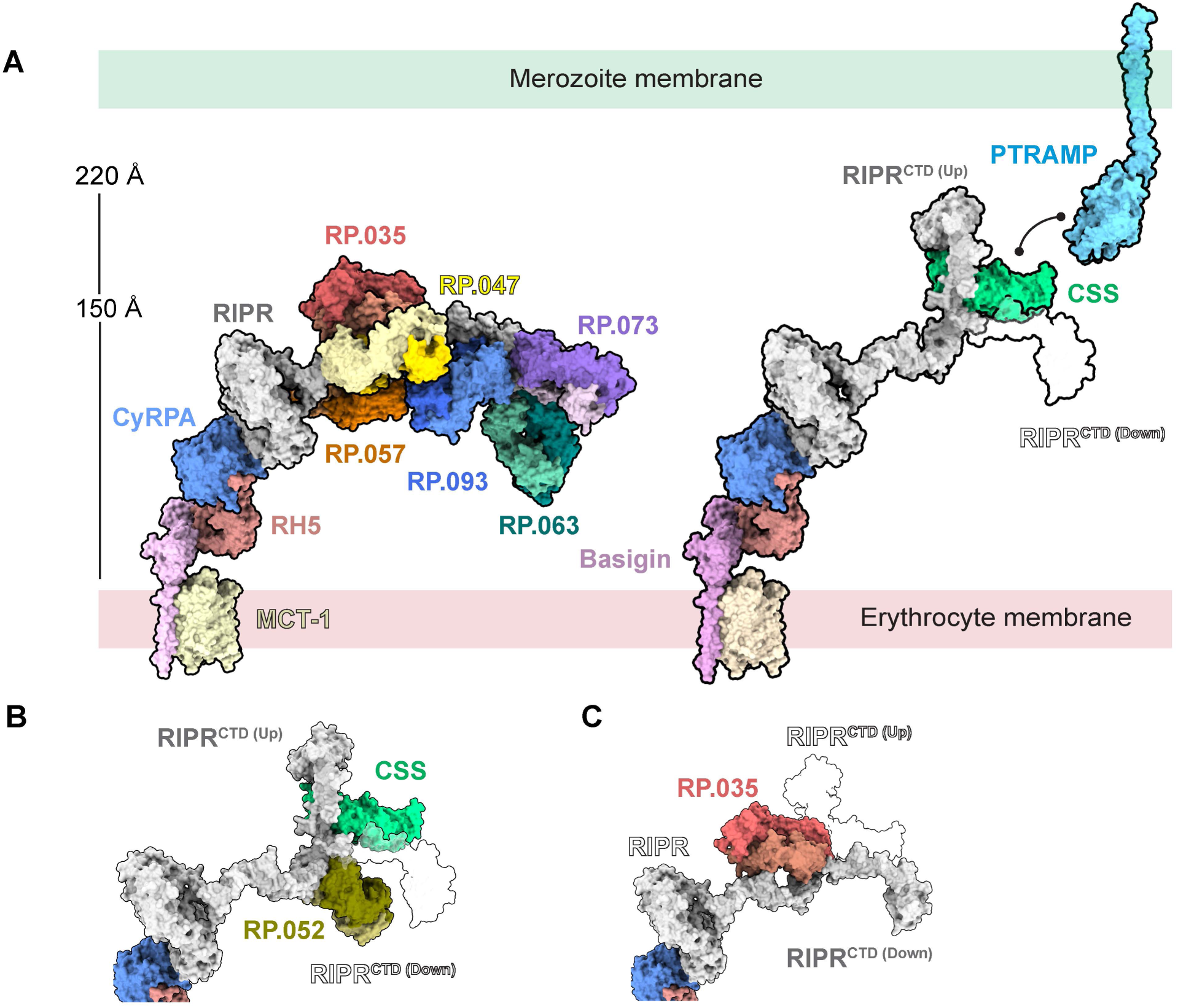
Anti-RIPR-Tail antibodies bound to the CSS-RIPR-CyRPA-RH5 complex. **A**) Structure of basigin:MCT1:RCR:RP.047:RP.057:RP.035:RP.093:RP.073:RP.063 complex (left) and basigin:MCT1:RCR:RP.092:RP.052:CSS complex with RP.092 and RP.052 omitted for clarity (right), both shown as a surface representation. The AlphaFold 2 model for PTRAMP (AlphaFoldDB: Q8I5M8) is also shown. The alternative RIPR^CTD (Down)^ conformation observed in the presence of the hexa-Fab complex is shown as a silhouette. **B**) Model of RP.052 bound to CSS:RIPR in the RIPR^CTD (Up)^ conformation, with the RIPR^CTD (Down)^ conformation shown as a silhouette. **C**) Model of RP.035 bound to RIPR in the RIPR^CTD (Down)^ conformation, with the RIPR^CTD (Up)^ conformation shown as a silhouette.

RIPR is also reported to bind the PC heterodimer through the RIPR^EGF (9-10)-CTD^ region.^14,35^ During sample preparation for cryo-EM, PC was complexed with RCR prior to the addition of the hexavalent antibody cocktail, however no cryo-EM density for PC was observed. We hypothesised one or more Fabs were interfering with the RIPR:CSS-PTRAMP interaction, and therefore sought to resolve this interaction using an alternative complex. Here, using a mixture of two anti-RIPR^EGF (7-8)^ antibodies, RP.092 and RP.052 (from Mix 1 in **Figure 2C**), we obtained a 4.0 Å reconstruction of these antibodies bound to RIPR^EGF (7-8)^ (PDB ID 9Q7C, EMD-72294) (**Figures S3D,E**). Low-resolution maps showed additional cryo-EM density consistent with CSS and the RIPR^EGF (9-10)-CTD^ region, but there was no density attributable to PTRAMP. This allowed us to rigid-body fit an AlphaFold 3 predicted model of CSS and RIPR^EGF (9-10)-CTD^. This yielded a composite model of the RIPR-Tail with CSS, which i) showed that CSS bound to the RIPR^EGF (9)^ region (**Figure S3F**) and ii) allowed us to superimpose this with our full-length RIPR model (**Figure 4A**). In this CSS-bound state, the conformation of the RIPR-Tail from RIPR^EGF (8)^ onwards was markedly different, with the RIPR^EGF (9-10)-CTD^ region shifting into a more upright conformation that runs perpendicular to the predicted merozoite membrane (which we termed RIPR^CTD (Up)^) at a distance of ∼220 Å from the erythrocyte membrane (**Figures 4A,B**).

These data were in contrast to the alternative conformation (RIPR^CTD (Down)^) that was observed in the presence of the hexavalent antibody cocktail. Here we noted that the anti-RIPR^EGF (8)^ antibody RP.035 would clash with RIPR^EGF (9)^ in the RIPR^CTD (Up)^ conformation (**Figure 4C**), suggesting that the binding of anti-RIPR Fabs in the RIPR^EGF (7-8)^ region could sterically modulate the conformation of the C-terminal region of the RIPR-Tail to RIPR^CTD (Down)^. In summary these data suggested that antibody binding along the RIPR-Tail could i) restrict flexibility (as evidenced by visualisation using cryo-EM), ii) modulate the RIPR^EGF (9-10)-CTD^ region into different conformations, and iii) potentially interfere with binding of CSS to the RIPR^EGF (9)^ region.

### Anti-RIPR^EGF (6-8)^ Fabs restrict RIPR-Tail flexibility in molecular dynamics simulations

In light of the structural data, we first hypothesised that the pools of anti-RIPR-Tail mAbs may mediate GIA by reducing RIPR-Tail flexibility. To assess this, we performed *in silico* molecular dynamics (MD) using our cryo-EM structures to explore the Fab-RIPR complexes in a dynamic system and to examine whether the binding of Fabs may be responsible for changes in the conformational ensemble of the RIPR-Tail. Given the number of systems to simulate, and our aim of assessing changes in flexibility of the RIPR-Tail ensemble, full-length RIPR protein and Fab regions were simulated with CALVADOS 3, a residue model for disordered and multi-domain proteins.^36,37^ Fabs were used for the model to reduce the computational requirement.

In line with its known flexibility, the simulations showed that RIPR-Tail takes up multiple conformations when not bound to Fabs (**Figure 5A,B, S4**). When the trivalent mix of anti-RIPR^EGF (9-10)-CTD^ Fabs (RP.063, RP.073 and RP.093) were docked onto RIPR, the simulations showed no difference in RIPR-Tail flexibility (**Figure 5C,D**). However, when anti-RIPR^EGF (6-8)^ Fabs were docked, the RIPR-Tail exhibited reduced flexibility. In the case of the trivalent mix of RP.035, RP.047 and RP.057 (“Mix 3”), the RIPR-Tail was typically held in an extended conformation away from the RIPR-Body, consistent with the RIPR^CTD (Down)^ cryo-EM structure observed with this combination of Fabs. In contrast, when we substituted RP.035 for RP.052, the RIPR-Tail was usually held in a compact conformation, with the CTD domain held closer to the RIPR-Body, again consistent with the RIPR^CTD (Up)^ conformation observed structurally with the RP.052 Fab (**Figure 5B,C**). We also found that, in these simulations, the RIPR-Tail extended rigidification was largely driven by the RP.035 Fab, with substantial additional constraint when further Fabs, such as RP.047 and RP.057 were included (**Figure 5E,F**). In a similar manner, the compaction of RIPR-Tail driven by RP.052 was enhanced when additional Fabs, such as RP.047 and RP.057, were added to the simulation (**Figure 5F,G**), suggesting synergistic interactions. These MD data were thus consistent with our finding that the pools of anti-RIPR^EGF (6-8)^ mAbs synergise with each other to mediate parasite growth inhibition (**Figure 2A,B**) and suggest constraint of RIPR-Tail flexibility as the underlying mechanism. Notably, our simulations indicated that further addition of the anti-RIPR^EGF (9-10)-CTD^ Fabs made no further impact on RIPR-Tail flexibility, suggesting a different mechanism accounts for their ability to synergise with the anti-RIPR^EGF (6-8)^ mAbs in the assays of GIA (**Figure 5H**).

**Figure 5.**
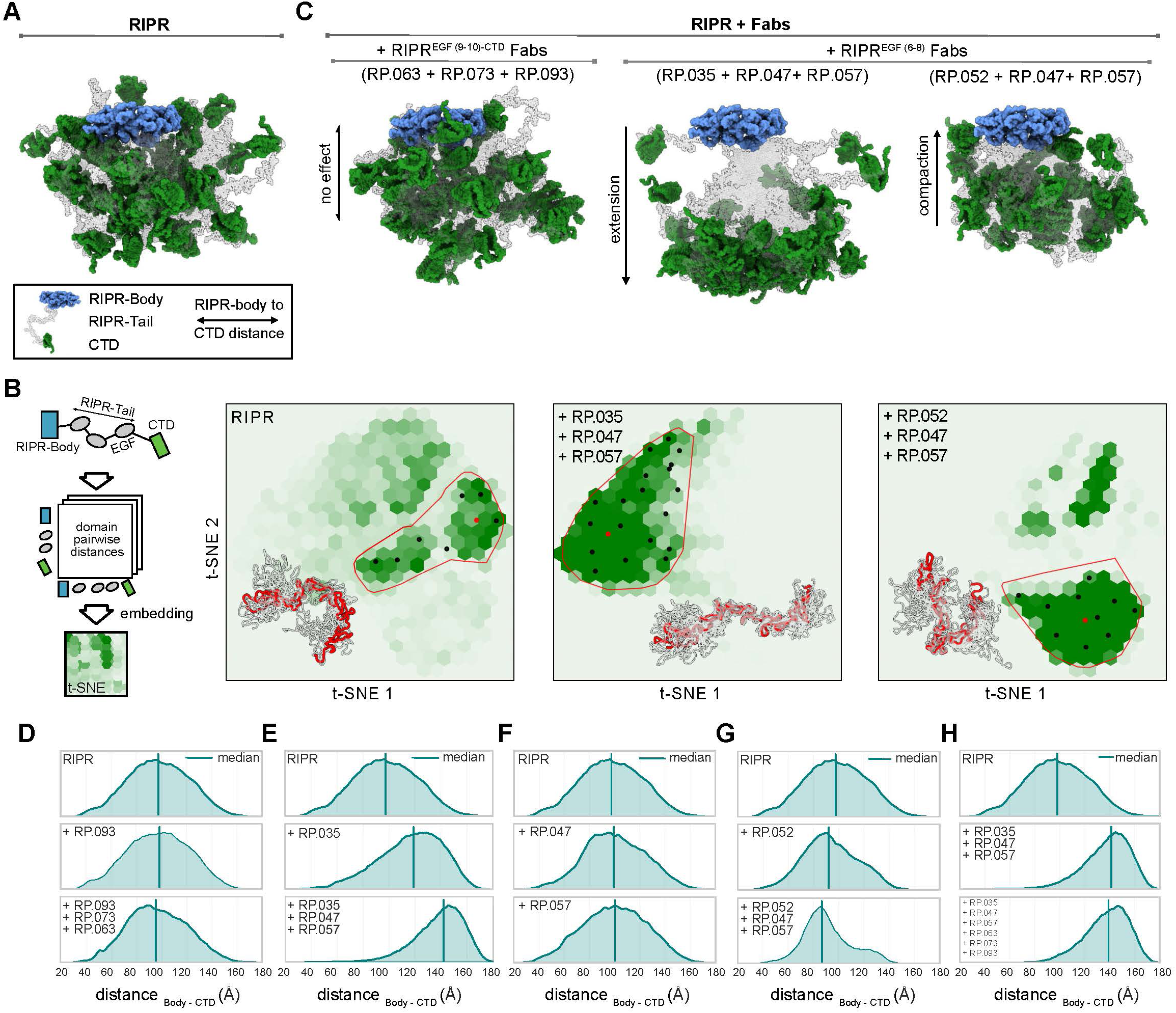
Computational analysis of RIPR conformational ensemble upon Fab binding using CALVADOS 3 molecular dynamics simulations. **A)** Cartoon illustrations of the RIPR conformational ensemble, as simulated with CALVADOS 3 for RIPR alone. The ensemble is shown with the N-terminal domain of RIPR (RIPR-Body) acting as an anchor, represented by a blue cartoon surface. The location variability of the RIPR^CTD^ region is represented by multiple green cartoon structures. The RIPR^EGF (5-10)^ region is illustrated as a transparent grey cartoon. **B**) Conformational space sampled by the RIPR-Tail during the MD simulation is shown in two-dimensional reduced space using t-distributed stochastic neighbour embedding (t-SNE). A schematic of the workflow used to generate the plots is shown on the left. Each panel shows the embedded space for RIPR alone or in complex with different Fabs. The largest structural cluster in each embedded space is outlined in red. Black dots mark representative ensemble structures for that cluster, with the centroid conformation highlighted in red. These representative structures from the cluster are reported as cartoon illustration in the bottom left corner, with the centroid coloured in red. **C**) Cartoon illustrations of the RIPR conformational ensemble in complex with different Fabs. RIPR is labelled as in (**A**). **D-H**) Distribution of the distance between the centre of mass (COM) of the RIPR-Body and RIPR^CTD^ region. In each plot, the solid line shows the median value of the distribution. The name of the target system is shown in the top left corner.

We further investigated the effects of the anti-RIPR^EGF (6-8)^ mAbs by studying the differences in the RIPR-Tail intra-molecular contacts in the simulations with and without individual Fabs. In the simulations with a single Fab, we found that all the differences in the RIPR-Tail intra-molecular contacts are located in regions corresponding to the interface with the Fab epitope as expected (**Figure S5A,B**). We then found that addition of the anti-RIPR^EGF (6-7)^ Fabs (RP.047 and RP.057) to the anti-RIPR^EGF^ ^(8)^ Fabs (RP.035 or RP.052) had a synergistic effect, with a large number of RIPR-Tail intra-molecular contacts appearing or disappearing when the Fabs were simulated together, but not when they were simulated separately (**Figure S5C,D**). If the Fabs did not behave synergistically to constrain the RIPR-Tail we would expect no additional contacts (aside from the Fab epitope interfaces) to be gained or lost when Fabs are simulated individually or together. These cooperative contacts had a lower magnitude for the combination of RP.047 and RP.057, suggesting that this effect is driven by the presence of RP.035 or RP.052. These data further supported our hypothesis that the synergistic GIA that we observed with the pools of anti-RIPR^EGF (6-8)^ mAbs is driven by constraining the flexibility of the RIPR-Tail.

### RIPR^EGF (9)^ binds PTRAMP-CSS with micromolar affinity

Our data suggested that the synergistic GIA mediated by the pools of anti-RIPR^EGF (6-8)^ mAbs is driven by these antibodies working in concert to constrain the flexibility of the RIPR-Tail. However, our MD simulations did not explain why constraining flexibility would cause GIA and, secondly, why pools of mAbs targeting RIPR^EGF^ ^(9-10)-CTD^ are GIA-negative but able to synergise with anti-RIPR^EGF^ ^(6-8)^ mAbs. Given our structural data showed RIPR^EGF^ ^(9)^ interacts with CSS, consistent with previous work that identified the PTRAMP-CSS heterodimer (PC) as a binding partner of RIPR^14^, we hypothesised that constraint of RIPR-Tail flexibility may affect this interaction and cause GIA. In order to assess this, we first characterised the interaction between recombinant RIPR and PC.

Recombinant PC was generated as described previously^27^ with the addition of an EndoH cleavage step to remove N-linked glycans. We verified glycan removal by gel shift assay and mass spectrometry (**Figure S6A,B, Data S4**). Binding affinities were then determined using SPR by chemically coupling either PC or RIPR-Tail to a CM5 chip. In line with previous reports analysing this interaction by SPR^27^, we observed low micromolar steady-state affinities for the interaction measured in both orientations (**Figure 6A,B**). We also then tested PTRAMP and CSS individually, and in agreement with Seager *et al.*^35^ and our structural data, we could detect RIPR-Tail binding to CSS but not to PTRAMP (**Figure S6C**). However, our measured affinities by SPR were much lower than those reported more recently by Seager *et al.* using biolayer interferometry (BLI) and a glycan-mutated PC construct.^35^ We therefore also tested our construct using BLI and capturing FL-RIPR using the anti-RIPR-Body mAb RP.122. Here we now measured a nanomolar affinity, but a 1:1 model could not be fitted to the data, rather a 2:1 heterogenous ligand model provided the best fit (**Figure S6D**). To try and reconcile the two different affinity measurements by SPR and BLI, we next performed mass photometry (MP) analysis of the PTRAMP-CSS-RIPR (PCR)-complex. We could only detect a low level of PCR-complex formation at both 100 nM and 250 nM concentrations, consistent with the low micromolar affinity of PC for RIPR as measured by SPR (**Figure 6C,D**). Given the complex model required to determine the kinetics, and heterogenous behaviour of the RIPR protein in BLI, we believed these data strongly suggest this interaction is low micromolar affinity. We thus further analysed the interaction by SPR and, finally, using the series of truncated RIPR-Tail proteins, confirmed that PC can bind a minimal truncation of RIPR^EGF (9-10)-CTD^ with low micromolar affinity, but not RIPR^EGF (10)-CTD^ or RIPR^CTD^ (**Figure 6E**). The necessity of RIPR^EGF (9)^ for CSS binding was consistent with the AlphaFold 3 prediction^35^ and our cryo-EM data.

**Figure 6.**
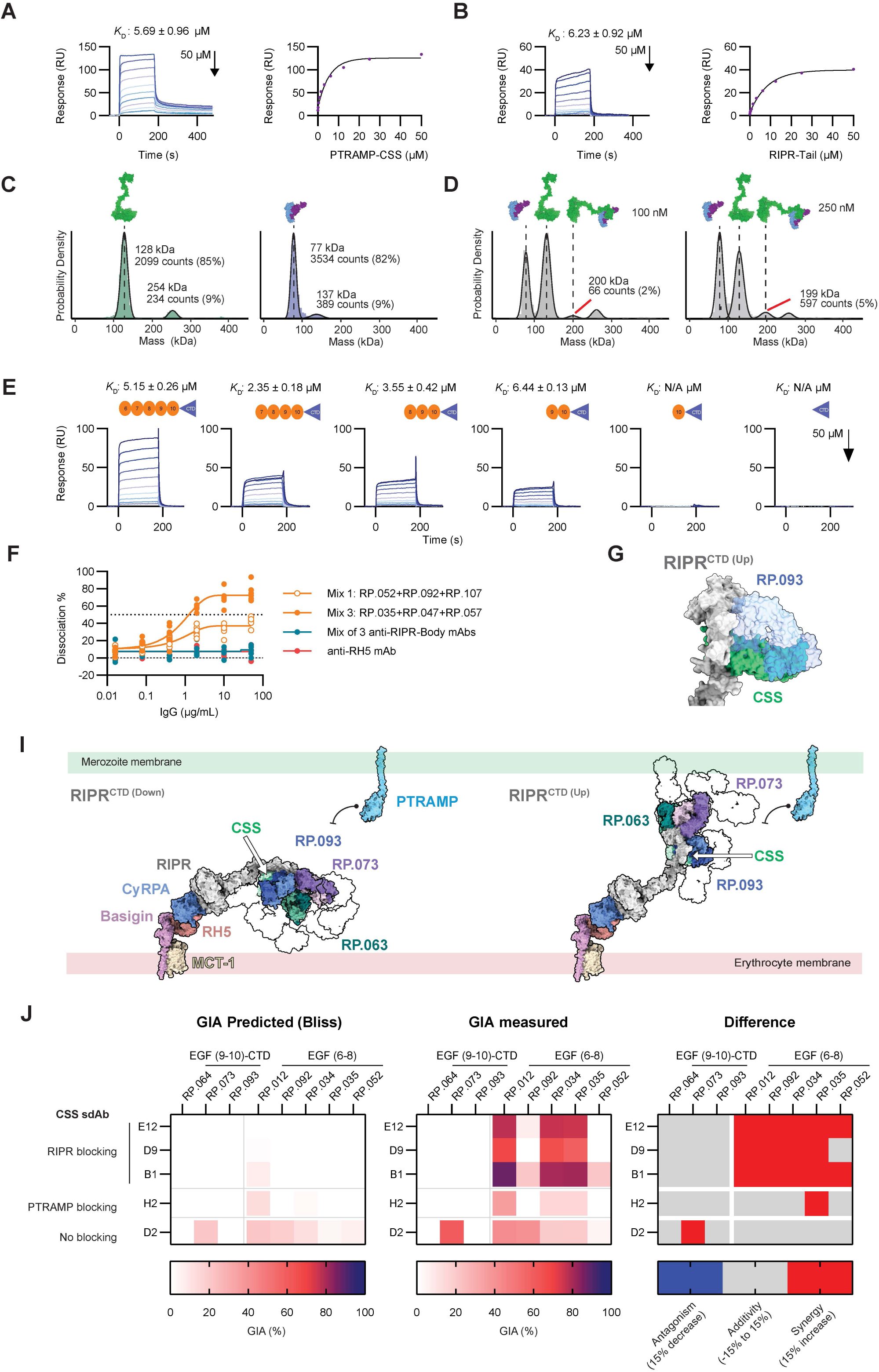
Anti-RIPR^EGF (6-8)^ mAbs disrupt the RIPR-PTRAMP-CSS complex. **A)** Left: Steady-state affinity, as assessed using SPR, of PTRAMP-CSS binding to RIPR-Tail coupled to the sensor surface. Sensorgrams are shown of a 10-step dilution curve beginning at 50 µM PTRAMP-CSS heterodimer. Right: Report points of the equilibrium binding levels of the 10-step 2-fold dilutions are fitted to the steady-state binding model for *K*_D_ determination. **B)** As for (A) but with PTRAMP-CSS heterodimer coupled to the sensor surface and RIPR-Tail used as the analyte. **C**) Mass distribution of RIPR binding to PTRAMP-CSS heterodimer after preincubation as determined by mass photometry showing RIPR alone or PTRAMP-CSS heterodimer alone, and **D**) 100nM or 250nM 1:1 mixture of RIPR and PTRAMP-CSS heterodimer. Histogram data are shown in green, purple, or grey, respectively. Gaussian curve fit shown as black line. **E**) Steady-state affinity, as assessed using SPR, of PTRAMP-CSS binding to indicated RIPR-Tail protein truncations coupled to the sensor surface. Sensorgrams are shown of a 10-step dilution curve beginning at 50 µM PTRAMP-CSS heterodimer. **F**) Dissociation ELISA where mixes of anti-RIPR antibodies were added in five-fold dilution series to the pre-formed FL-RIPR: PTRAMP-CSS complex. Resulting biotinylated PTRAMP-CSS was measured and the percent dissociation calculated. **G**) Model of RP.093 bound to RIPR superimposed onto model of CSS:RIPR in the RIPR^CTD (Up)^ conformation. **I)** Composite model of RCR:RP.093:RP.073:RP.063 complex (as in Figure 4A) in RIPR^CTD^ ^(Down)^ (left) and RIPR^CTD (up)^ (right) conformation shown as a surface representation. Fabs structures are indicated by colour with the rest of the IgG modelled as silhouettes (PDB ID: 1IGT). CSS model (green) is docked onto RIPR as indicated by white arrow. The AlphaFold 2 model for membrane PTRAMP (AlphaFoldDB: Q8I5M8) is also shown. **J**) Heatmap of anti-CSS sdAb and anti-RIPR mAb combination GIA analysis. The indicted antibodies were tested in pairwise combinations at 1mg/mL per mAb/sdAb. Left: predicted % GIA (Bliss additivity) of each combination; centre: measured % GIA of each combination; right: predicted % GIA subtracted from the measured % GIA. ≥15% GIA over predicted was classified as synergistic (red), ≤15%; less than predicted as antagonistic (blue), otherwise the result was classified as additive (grey). GIA data shown in **Figure S6H**.

### Anti-RIPR^EGF (6-8)^ mAbs dissociate the RIPR:PTRAMP-CSS complex and enable antibody blockade of the RIPR-CSS interaction

The PCRCR-complex is reported to pre-form within the parasite^14^, meaning the RIPR-CSS binding interface would not be accessible to antibody on the merozoite surface. This could explain why anti-RIPR^EGF (9-10)-CTD^ mAbs are GIA-negative when tested alone. In light of our data, we hypothesised that constraint of RIPR-Tail flexibility, caused by binding of anti-RIPR^EGF (6-8)^ mAbs, disrupts this low-affinity interaction and exposes epitopes in the RIPR-CSS binding region of the pentameric complex. Having characterised reagents to assess the interaction, we next developed a PCR-complex dissociation ELISA to measure the ability of anti-RIPR^EGF (6-8)^ mAbs to induce PC dissociation from RIPR. Here, FL-RIPR was pre-incubated with biotinylated PC prior to addition of mAbs and measurement of bound PC. Addition of control anti-RH5 mAb or pools of anti-RIPR-Body mAbs showed no effect on PC binding. In contrast, when using Mix 1 and Mix 3 of the anti-RIPR^EGF (6-8)^ mAbs, 40-80% dissociation of PC from FL-RIPR was observed which titrated out in the assay as mAb concentration was reduced (**Figure 6F**).

Partial dissociation of the recombinant PCR-complex in the presence of anti-RIPR^EGF (6-8)^ mAbs was consistent with partial *in vitro* GIA observed with the same mixes of antibodies (**Figure 2C**). Given anti-RIPR^EGF (9-10)-CTD^ antibodies can then potentiate this GIA, we next hypothesised that these antibodies prevent the PCR-complex re-forming. We first investigated whether any anti-RIPR mAbs could block the binding of RIPR to PC. As positive controls, we produced five previously reported anti-CSS sdAbs^14^ (**Figure S6E**) that either block the RIPR-CSS interaction (B1, D9, E12), block the CSS-PTRAMP interaction (H2), or that do not block either interaction but show GIA activity (D2). Binding of the sdAbs to PTRAMP-CSS was assessed by ELISA (**Figure S6F**). Four sdAbs behaved as expected, however, D2 showed low reactivity likely due to removal of the N-linked glycan at residue N88 on our recombinant CSS which is involved in D2 binding.^14^ We next used a competition ELISA to screen sdAb controls and the anti-RIPR mAb panel for blockade of PC binding to RIPR (**Figure S6G**). As expected, the D9 sdAb completely blocked the PC-RIPR interaction, whereas D2 did not. Notably, we only identified two mAbs, RP.064 and RP.093, that partially blocked formation of the PCR-complex when tested alone. These data for RP.093 were consistent with our structural observations showing that the variable region of this Fab sterically occluded the binding site of CSS when bound to RIPR (**Figure 6G**). Moreover, the inability of the wider panel of anti-RIPR^EGF (9-10)-CTD^ mAbs, when tested alone, to show consistent blockade of the PC-RIPR interaction was consistent with their variable ability to potentiate GIA of the anti-RIPR^EGF (6-8)^ mAbs when tested in pair-wise combinations (**Figure 2A,B**). We thus instead sought to assess the two trivalent mixtures of anti-RIPR^EGF (9-10)-CTD^ mAbs (Mixes 1/3 and 2 from **Figure 2C**) for blockade of the RIPR-PC interaction, given their consistent ability to synergise in the GIA assay. However, we were unable to develop a reliable *in vitro* assay to assess the blocking ability of the trivalent mAb mixtures by ELISA, SPR or MP (data not shown). We therefore structurally modelled the three anti-RIPR^EGF (9-10)-CTD^ mAbs (Mix 1/3) bound to both the RIPR^CTD(Down)^ and RIPR^CTD(Up)^ conformations. In both scenarios, the bound mAbs would sterically hinder the re-binding of a membrane-bound PTRAMP-CSS complex to the RIPR-Tail (**Figure 6I**).

Finally, we also hypothesised that if blockade of RIPR-PC re-binding is associated with GIA following constraint of the RIPR-Tail and disruption of the RIPR-PC interaction by anti-RIPR^EGF (6-8)^ mAbs, then the same GIA assay outcomes should be observed when combining these mAbs with anti-CSS antibodies that similarly block the RIPR-CSS binding interface. We thus tested the anti-CSS sdAbs in combination with five anti-RIPR^EGF (6-8)^ mAbs and three anti-RIPR^EGF (9-10)-CTD^ mAbs. Consistent with reported data^14^, there was some modest GIA with D2 but otherwise no GIA observed with the anti-CSS sdAbs alone, including the three clones that block the RIPR-CSS interaction (B1, D9, E12). In contrast, synergistic GIA was observed when these three anti-CSS sdAbs were combined with the anti-RIPR^EGF (6-8)^ mAbs but not with the anti-RIPR^EGF (9-10)-CTD^ mAbs (**Figure 6J, S6H**). The measured synergy between the anti-RIPR^EGF (6-8)^ mAbs and anti-CSS sdAbs was consistent with the level of synergy seen with the anti-RIPR^EGF (6-8)^ mAbs and anti-RIPREGF (9-10)-CTD mAbs (**Figure 2A,B, S2A**) The same effects were not observed with the two sdAbs that do not block the RIPR-CSS interaction. These data strongly supported our hypothesis that mAb binding to the RIPR^EGF (6-8)^ region of the RIPR-Tail, distal to the CSS binding site, leads to epitope accessibility on the RIPR-CSS binding interface and that blockade of this interaction is associated with synergistic inhibition of parasite growth.

### Enhancing GIA of the R78C clinical vaccine candidate

A RIPR-CyRPA candidate vaccine, called R78C, is currently in Phase 1 clinical trials, and contains the RIPR^EGF (7-8)^ region of RIPR fused to CyRPA.^17^ Our data here suggested that the addition of RIPR^EGF (6)^ and RIPR^(9-10)-CTD^ could improve this vaccine. To assess this hypothesis, we purified a pool of total IgG from healthy UK adults vaccinated with the R78C/Matrix-M vaccine candidate, this showed ∼78% GIA when tested alone at 6 mg/mL total IgG (**Figure 7A**). This GIA was reversed to ∼20% in the presence of 5μM CyRPA protein, and completely reversed in the presence of 5μM CyRPA and RIPR proteins, confirming antigen-specificity of the vaccine-induced GIA. In parallel we identified a pool of four GIA-negative but synergistic anti-RIPR ^(9-10)-CTD^ mAbs (RP.063, RP.064, RP.073 and RP.085) and a second pool of two anti-RIPR^EGF (6)^ mAbs (RP.047 and RP.107). Neither pool showed GIA when tested alone but ∼30% GIA when combined (**Figure 7A**), in line with our previous data.

**Figure 7.**
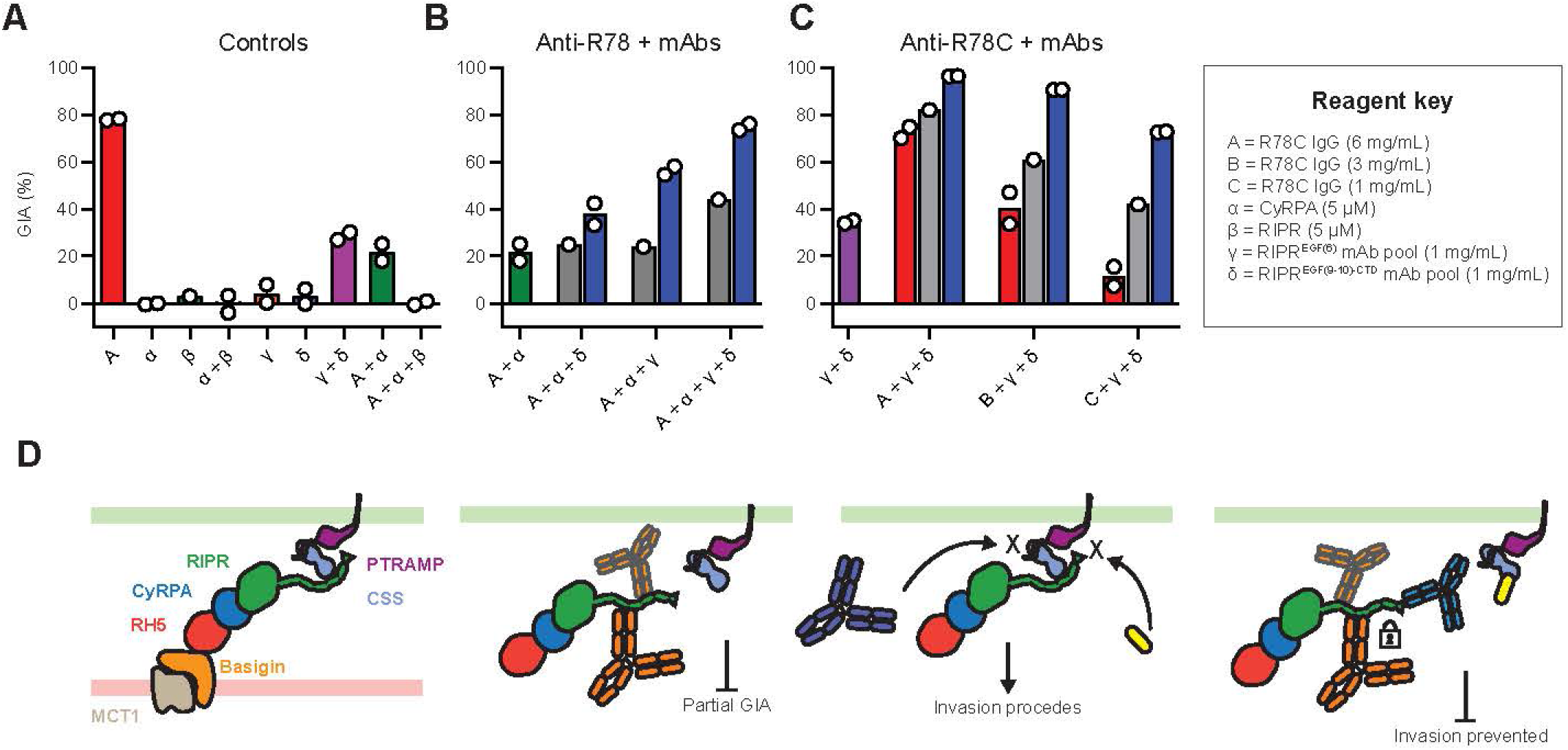
Anti-RIPR-Tail mAbs improve GIA of the R78C clinical vaccine candidate. **A)** *In vitro* GIA against 3D7 clone *P. falciparum* using i) a pool of purified total IgG from R78C/Matrix-M healthy UK adult vaccinees tested at 6 mg/mL; ii) protein alone controls (CyRPA and RIPR) both at 5 µM; iii) a pool of anti-RIPR^EGF (9-10)-CTD^ mAbs (RP.063, RP.064, RP.073 and RP.085) and a pool of anti-RIPR^EGF (6)^ mAbs (RP.047 and RP.107), both at 1 mg/mL; and iv) combinations of test samples. **B**) GIA of the purified total IgG from R78C vaccinees combined with CyRPA protein along with addition of either or both of the anti-RIPR mAb pools (all concentrations as in panel A). Grey bars show % GIA predicted by Bliss additivity (PBA) of the test combinations, and blue bars how the measured % GIA. **C**) GIA of the purified total IgG from R78C vaccinees (without CyRPA protein) tested at 6, 3 or 1 mg/mL alone (red bars) or with addition of both of the anti-RIPR mAb pools (at a fixed total concentration of 2 mg/mL). Grey and blue bars indicate predicted and measured GIA as in panel B. Bars show mean, N = 2 independent replicates. **D**) Cartoon summary of the proposed two-step mechanism for GIA mediated by anti-RIPR-Tail antibodies. left: the full *P. falciparum* PCRCR-complex binds to basigin-MCT1 complexes on the erythrocyte surface. Centre left: binding of anti-RIPR^EGF (6-8)^ mAbs (orange) can lead to partial GIA. Centre right: anti-RIPR ^(9-10)-CTD^ mAbs (blue) or anti-CSS sdAbs (yellow) that block the RIPR-CSS binding interface are normally unable to access their epitopes and so invasion proceeds (no GIA). Right: Binding of anti-RIPR^EGF (6-8)^ antibodies constrains RIPR-Tail conformational flexibility and disrupts the low affinity RIPR-CSS interaction. Exposure of the RIPR-CSS binding interface now enables binding of anti-RIPR ^(9-10)-CTD^ mAbs or anti-CSS sdAbs to their epitopes, thus blocking further RIPR-CSS interaction and synergistically enhancing GIA.

We next tested the R78C IgG combined with 5μM CyRPA protein to assess only the GIA attributable to the anti-RIPR^EGF (7-8)^ polyclonal IgG (i.e. the R78 portion of the vaccine). Addition of both the anti-RIPR ^(9-10)-CTD^ mAb pool and the anti-RIPR^EGF (6)^ mAb pool improved GIA above the predicted additive, but the largest synergistic increase was seen to 76% GIA when both pools were added (as compared to the predicted additive of ∼40% GIA) (**Figure 7B**). These data confirmed the mAb pools were capable of synergising with vaccinated human anti-RIPR^EGF (7-8)^ polyclonal IgG. Finally, we repeated this experiment without addition of 5μM CyRPA protein to assess outcome on the total anti-R78C response. Addition of both mAb pools showed a synergistic effect, reaching >90% GIA when combining the pools of mAbs with 6 mg/mL total IgG (**Figure 7C**). Titration of the anti-R78C IgG to 3 mg/mL and 1mg/mL in the assay further demonstrated the synergistic effect of the mAb pools with overall GIA levels maintained well above the predicted additive. These data indicate that anti-RIPR antibody synergism still occurs in the presence of anti-CyRPA human IgG, and provide a framework to guide re-design of next-generation vaccines that include more domains from the RIPR-Tail.

## DISCUSSION

The RIPR protein is highly conserved across the *Plasmodium* genus, and forms part of the essential PCRCR-complex used by *P. falciparum* parasites to invade erythrocytes.^11,14,29^ Conditional gene knockouts of any PCRCR protein show merozoites that can deform the host cell membrane but which fail to internalise.^14,15^ Previous cryo-EM data showed that RIPR consists of an ordered, multi-domain core (RIPR-Body) which spans from the N-terminus to EGF-like domain 4 and which binds CyRPA (which in turn binds RH5). Subsequently, from EGF-like domain 5 onwards, the RIPR molecule extends into a flexible elongated tail, whose structure could not be resolved, but which was shown to bind the PTRAMP-CSS complex on the merozoite surface; thereby bridging the erythrocyte and parasite membranes during invasion.^27^ Understanding how antibodies against this invasion complex can mediate functional growth inhibition of *P. falciparum* parasites has become central to next-generation blood-stage malaria vaccine development strategies.

Previous analyses of sera from FL-RIPR-vaccinated animals reported that functional IgG are directed against EGF-like domains 5-8 in the RIPR-Tail.^17^ Consistent with this, vaccination of animals with constructs spanning RIPR^EGF (5-8)^ can raise polyclonal antibody responses that show functional GIA against *Plasmodium* parasites.^11,13,17,38^ Here, we expanded on these observations and identified that polyclonal anti-FL-RIPR IgG also contains GIA-negative antibodies that i) bind within the RIPR^EGF (8-10)-CTD^ region of the RIPR-Tail and ii) potentiate the GIA-positive anti-RIPR^EGF (5-8)^ IgG.

To understand this, we isolated 83 unique anti-RIPR mAbs from the humanised Kymouse platform, of which 67 bound the RIPR-Tail. Intriguingly no mAb showed GIA when tested in isolation, despite the vaccine-induced polyclonal purified total IgG showing high-level GIA and the mAbs binding multiple epitopes across the RIPR-Tail. To rationalise this, we set up a comprehensive pairwise GIA screen using the anti-RIPR-Tail mAb panel. This identified two zones of epitopes along the RIPR-Tail, with a “neutralisation zone” comprising RIPR^EGF (6-8)^ and a “synergy zone” comprising RIPR^EGF (9-10)-CTD^. Our data enabled us to propose a model for how maximum GIA is achieved via a two-step synergistic mechanism that occurs as antibodies bind along the length of the RIPR-Tail (**Figure 7D**). Here, anti-RIPR^EGF (6-8)^, but not anti-RIPR^EGF (9-10)-CTD^, antibodies can synergise to constrain RIPR-Tail conformational flexibility, which is consistent with their respective abilities to mediate GIA when tested alone. MD simulations indicate that the RIPR-Tail can take up multiple conformations, but that binding of anti-RIPR^EGF (6-8)^ mAbs rigidifies the molecule, either by holding the RIPR-Tail in an extended conformation, as driven by mAb RP.035, or by compacting the RIPR-Tail as driven by mAb RP.052. Addition of further Fabs potentiated this effect thus providing a mechanistic basis for the synergistic GIA observed with anti-RIPR^EGF (6-8)^ antibodies.

We also observed that binding of anti-RIPR^EGF (6-8)^ IgG can partially displace recombinant PC heterodimer bound to FL-RIPR. Consistent with another report^35^, we showed that EGF-like domain 9 is necessary for binding of *P. falciparum* RIPR to CSS, with SPR and MP data indicating this interaction to be low micromolar affinity. We hypothesise that disruption of this weak RIPR-CSS interaction by anti-RIPR^EGF (6-8)^ IgG is directly linked to the reduction in RIPR-Tail conformational flexibility, and that in turn this disruption of the pentameric PCRCR-complex scaffold inhibits parasite invasion of the erythrocyte. In support of this, we also structurally defined two RIPR-Tail conformations, with data suggesting that the binding of anti-RIPR Fabs in the RIPR^EGF (7-8)^ region can sterically modulate the RIPR^EGF (9-10)-CTD^ region. We propose this modulation of RIPR-Tail ultimately interferes with its ability to remain bound to CSS, however, a direct link between these observations remains to be formally demonstrated.

It has also been reported that the PCRCR-complex pre-forms within the parasite, prior to its release to the parasite surface.^14^ Consequently, antibodies that block interactions between the complex components, such as RH5-CyRPA, RIPR-CSS or CSS-PTRAMP, are consistently GIA-negative because their epitopes are not surface-exposed.^14,17,22,26^ Dissociation of the membrane-bound PTRAMP-CSS heterodimer from RIPR, caused by binding of anti-RIPR^EGF (6-8)^ IgG, would in contrast expose epitopes present at the RIPR-CSS binding interface and allow for binding and potential functional activity. In support of this, our data showed that anti-RIPR^EGF (9-10)-CTD^ IgG, or anti-CSS sdAbs that directly block the RIPR-CSS interaction, are GIA-negative when tested alone, however, they potentiate the GIA of anti-RIPR^EGF (6-8)^ IgG when combined. Indeed, our structural data suggested that mAbs bound to the C-terminal region of RIPR would directly or sterically hinder the re-binding of a membrane-bound PTRAMP-CSS complex to the RIPR-Tail. Similarly, we also tested a panel of anti-CSS sdAbs^14^, and only those that directly block the RIPR-CSS interaction synergised with the anti-RIPR^EGF (6-8)^ IgG. These data thus strongly suggest that prevention of PCRCR-complex re-association (via RIPR-Tail rebinding to CSS) at the parasite surface is a contributing factor to the overall maximal GIA we observed. These data also suggest that GIA-positive nanobodies reported to bind other epitopes on CSS or PTRAMP^14^ likely exert their functional activity by other, as yet undefined, mechanisms. Moreover, the need for multiple mechanisms to achieve high-level GIA likely explains why no single high-potency anti-RIPR mAb has been reported. Whether a single multi-specific antibody-like molecule could be designed to concurrently constrain RIPR-Tail flexibility and block the RIPR-CSS binding site is a question worthy of future investigation.

Understanding whether polyclonal antibody responses targeting antigens in the PCRCR-complex can synergise has been an active area of investigation.^10,22,32,39,40^ One study has shown that a class of GIA-negative anti-RH5 antibody can slow down the rate of merozoite invasion, thereby potentiating growth inhibitory antibodies by providing more time for them to bind their target epitopes^22^. Another study showed that antibodies binding adjacent epitopes on CyRPA can form lateral heterotypic interactions, thereby enhancing binding and improving GIA.^26^ Our studies on RIPR expand on these reports and identify new mechanisms that explain how GIA-negative antibody clones can synergise to exert high-level GIA. Our results also raise interesting questions for next-generation blood-stage malaria vaccine design. Some structure-based vaccine design strategies for malaria have focused on designing immunogens that encode potent neutralising antibody epitopes.^18,41,42^ RIPR, however, may not be amenable to this approach due to the lack of single potent epitopes.

Instead, our data suggest an alternative approach targeting a range of RIPR domains may be more effective. This may also help guard against vaccine escape by spreading immune pressure over a larger target antigen area, rather than focussing on single epitopes. In line with this, our experiments using R78C human vaccinee IgG show that the addition of the RIPR^EGF^ ^(6)^ and/or RIPR^EGF^ ^(9-10)-CTD^ domains could substantially improve the potency of this vaccine candidate. This work remains in progress using the data reported here as a blueprint for next-generation RIPR-based vaccine designs.

## LIMITATIONS OF THE STUDY

A limitation of this work is that our antibody panel was isolated from the humanised Kymouse platform, while the target population for a RIPR-based blood-stage malaria vaccine is African children. Notably a few mAb clones, isolated from wild-type mice, have shown modest GIA when used alone^17,28^, in contrast to the data reported here where all clones were GIA-negative. Consistent with our data, these murine mAbs all bind within the RIPR^EGF (5-8)^ region, but this does suggest some species-dependence in the vaccine-induced anti-RIPR antibody repertoire. Importantly we also show here that the polyclonal anti-RIPR^EGF (7-8)^ response induced in human R78C vaccinees was GIA-positive, consistent with our observations using the pools of mAbs isolated from humanised mice. Nonetheless, future work will need to confirm the generalisability of our results to the human population and in particular African children. Secondly, our study also only investigated mAb function and potency against 3D7 clone *P. falciparum* using the *in vitro* assay of GIA, and we did not investigate impact of RIPR polymorphism. Nonetheless, we have previously shown that GIA-positive antibodies confer protection in humanised mice and *Aotus* monkeys challenged with *P. falciparum*^21,22,43,44^, and that a vaccine inducing high-level GIA against RH5 protects against clinical malaria in young African children^9^. Parasite sequencing data have also shown the RIPR-Tail is highly conserved, with only 3 SNPs across this 370 amino acid region (A755G, Y985N and I1039M) reported at >5% frequency globally.^45-47^ Our data will now facilitate a larger-scale analysis of anti-RIPR mAbs against field isolates to assess for any variation in functional antibody potency that could be driven by parasite genetic variation.

## AUTHOR CONTRIBUTIONS

Conceived and performed experiments and/or analysed the data: BGW, JRB, JBS, CAR, MC, DQ, KM, AH, SAB, SSRR, AR, LB, BB, SW, NM, LDWK, FRD, CAG, CG, FB, CMD, STR, ABW, SJD.

Contributed reagents, materials, and analysis tools: MC, SES, JS, AMM, WBS, CMD, ABW. Performed project management: JP, SS, KS.

Wrote the paper: BGW, JRB, SJD.

## ACKNOWLEDGEMENTS

The authors are grateful for the assistance of Amy Boyd, Jenny Bryant, Patries Fisher, Carolyn Nielsen, Lana Strmecki, Mariola Zaleska, David Staunton (University of Oxford); Kazutoyo Miura (NIAID, NIH); Paul Kellam, Sara Chalk, Lauren Weatherill, Emma MacGregor (Kymab/Sanofi); Jacqueline Kirchner, Holger Kanzler, Hedda Wardemann, Annie Zumsteg, Laine Milburn, Deborah Roby (Gates Foundation); Letitia Jean for arranging contracts (University of Oxford); We thank Novavax for provision of Matrix-M® adjuvant in the VAC089 clinical trial, and the VAC089 trial participants.

## INCLUSION AND DIVERSITY

We support inclusive, diverse, and equitable conduct of research.

## SUPPLEMENTARY TABLE AND FIGURE LEGENDS

**Figure S1.**
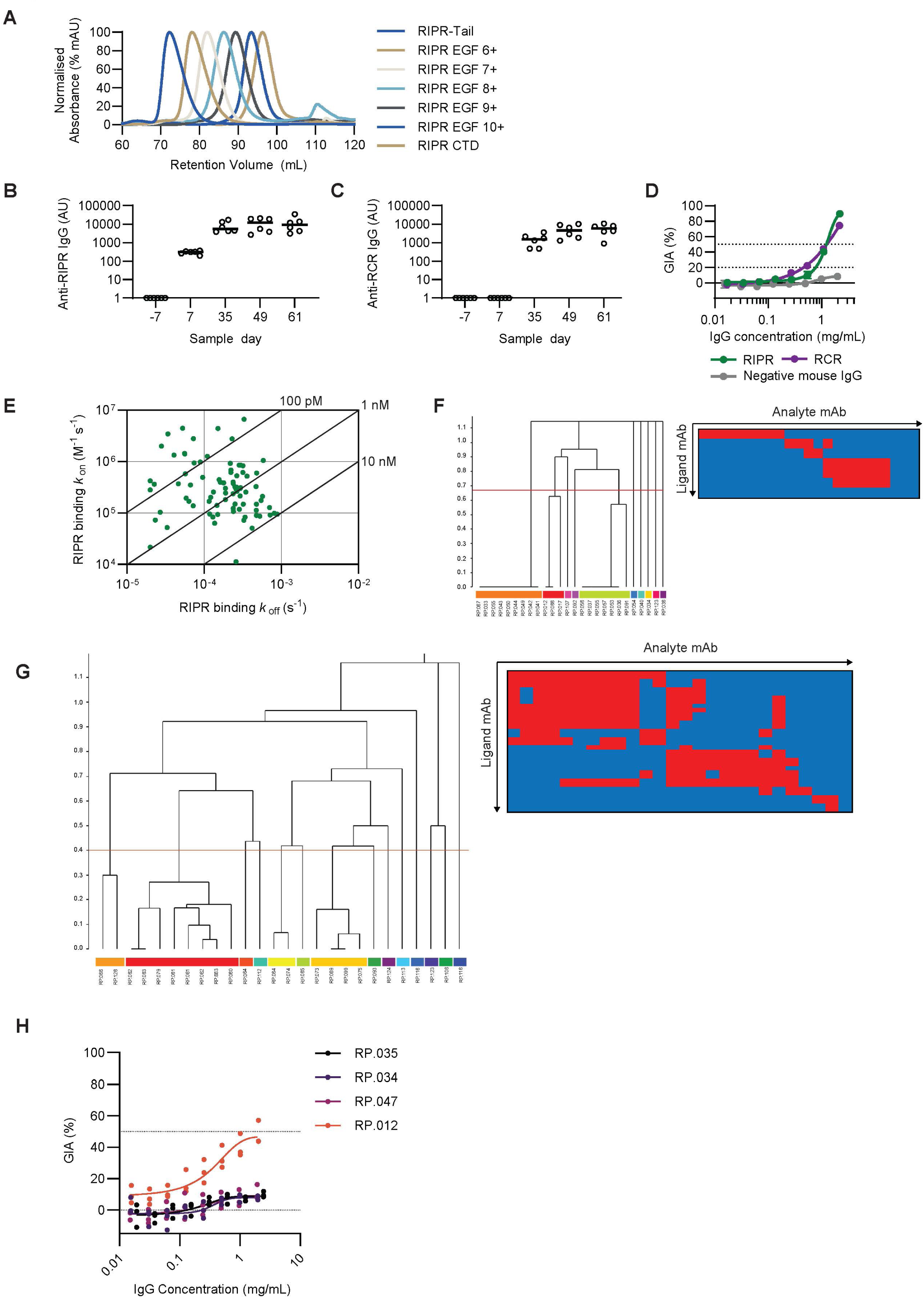
Development and evaluation of anti-FL-RIPR antibodies; related to Figure 1. **A)** Size exclusion chromatograms overlaid for each of the RIPR-Tail protein truncations. **B**) Anti-RIPR serum IgG ELISA titres shown for samples taken on days -7, 7, 35, 49 and 61 for Kymice immunised with FL-RIPR, reported in arbitrary units (AU). Individual and median group responses are shown. **C**) Anti-RCR-complex serum IgG ELISA titres shown for samples taken on days -7, 7, 35, 49 and 61 for animals immunised with the RCR-complex, reported in AU. Individual and median group responses are shown. **D**) Single-cycle *in vitro* GIA assay using 3D7 clone *P. falciparum* parasites. Total IgG was purified from day 61 vaccinated Kymouse serum samples and titrated in the GIA assay. Serum from N=3 animals were pooled for each group. Each point represents the mean of three technical replicates. Negative mouse IgG came from serum of non-vaccinated Kymice. **E**) Iso-affinity plot showing kinetic rate constants for binding of mAbs to FL-RIPR as determined by HT-SPR. Diagonal lines represent equal affinity (*K*_D_) = *k_off_*/*k_on_*. **F**) Left, competition profiles were used to cluster anti-RIPR^EGF (5-8)^ mAbs into epitope communities represented by coloured regions of the dendrogram using the McQuitty clustering method in the Carterra Epitope software. Right, heatmap displaying competition interactions between anti-RIPR^EGF (5-8)^ mAbs as ligands (y axis) and analytes (x axis). Red squares indicate competition between the mAb pair, and blue squares indicate no competition between the mAb pair. **G**) As for (F) but with anti-RIPR^EGF (9-10)-CTD^ mAbs. **H**) GIA assay as per (D). mAbs were titrated from 2 mg/mL in a two-fold dilution series. N=3 independent repeats, each data set is fitted with a Richard’s five-parameter dose-response curve with no constraints.

**Figure S2.**
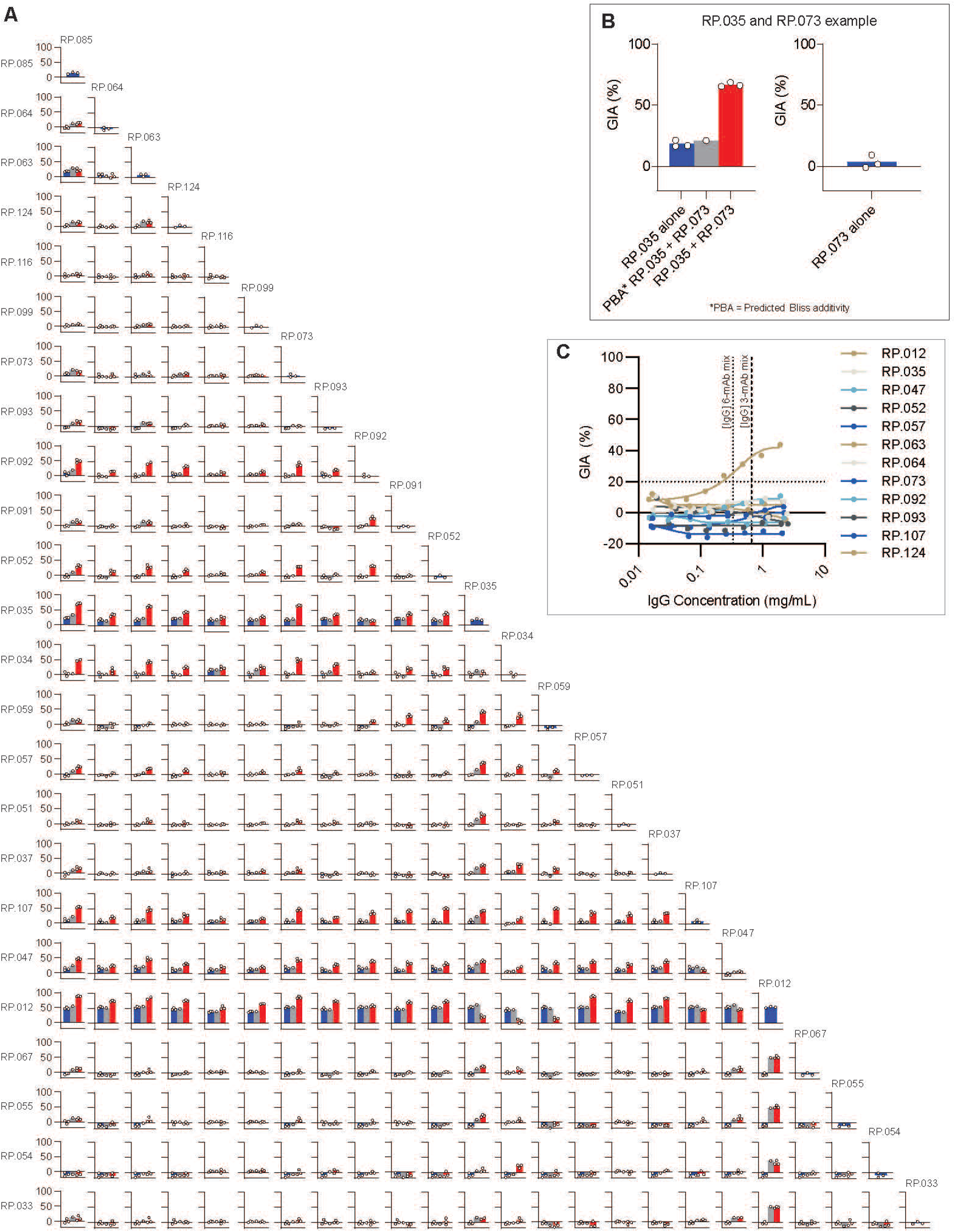
Anti-FL-RIPR mAb GIA synergy screen; related to Figure 2. **A)** Each mAb was tested alone (at 1 mg/mL) or in combination with a second mAb (at 1 mg/mL each). Blue bar represents the test mAb alone, the grey bar shows the predicted GIA (assuming Bliss additivity) and the red bar shows the measured result. Mean values and individual technical replicates shown. **B**) Example inset shows plots from (A) for mAb RP.035 combined with RP.073 expanded for clarity. **C**) Single-cycle *in vitro* GIA assay using 3D7 clone *P. falciparum* parasites. Anti-RIPR mAbs were titrated from 2 mg/mL in a two-fold dilution series. X-axis line at 0.66 mg/mL (dashed) represents the highest concentration of each mAb in the 3-mAb mixes tested in Figure 2C whilst the dotted line at 0.33 mg/mL the concentration of each mAb in the 6-mAb mixes tested in Figure 2C. N=1 assay with 3 technical repeats; each data set is fitted with a Richard’s five-parameter dose-response curve with no constraints.

**Figure S3.**
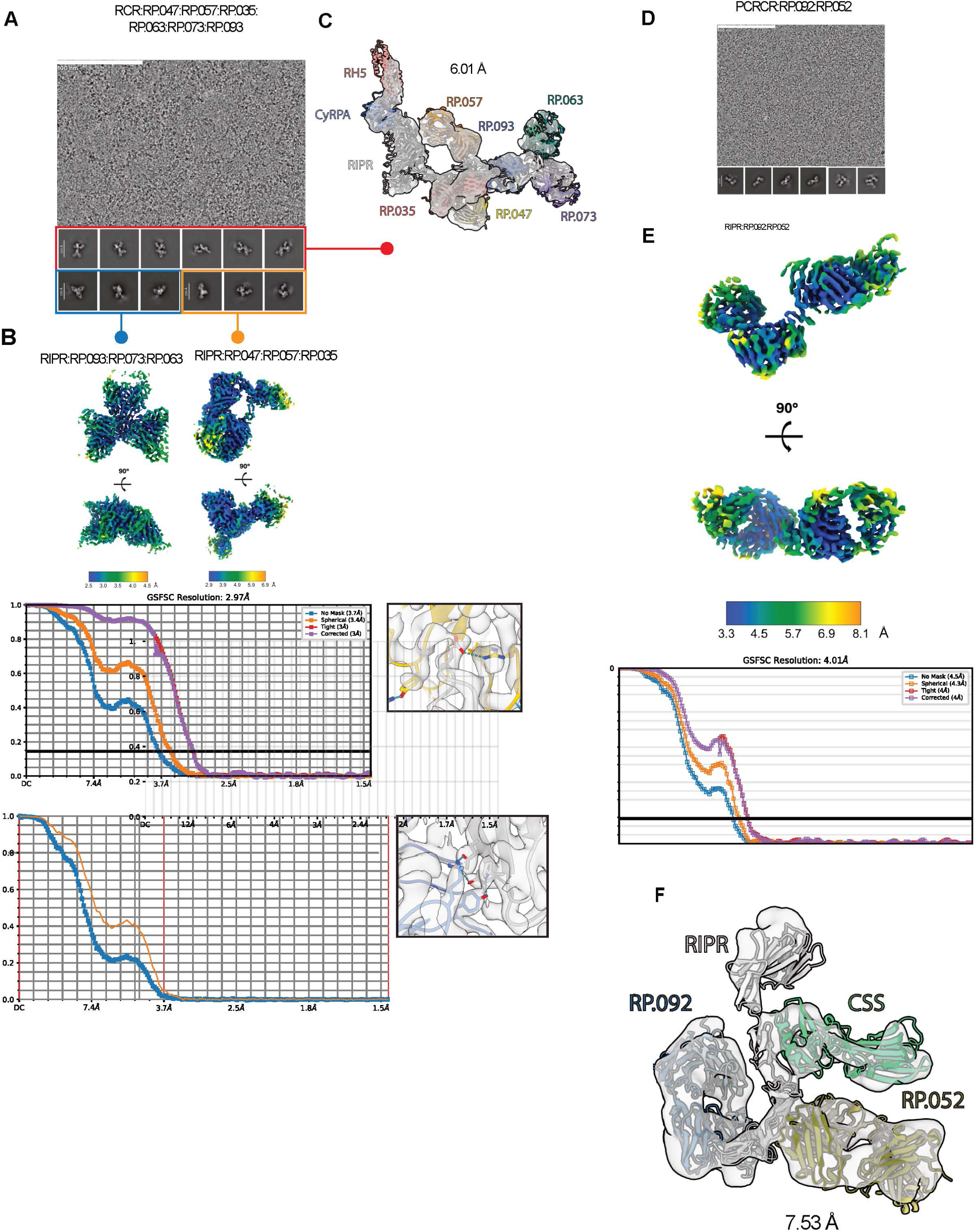
Building the full-length composite model of RCR:RP.047:RP.057:RP.035:RP.063:RP.073:RP.093; related to Figure 3 and Structural data of PCRCR:RP.092:RP052; related to Figure 4. **A**) Representative micrograph and 2D classes of the RCR:RP.047:RP.057:RP.035:RP.063:RP.073:RP.093 complex sample from which the maps and models for RIPR:RP.093:RP.073:RP.063 (PDB ID 9Q6B, EMD-72265); RIPR:RP.047:RP.057:RP.035 (PDB ID: 9Q69, EMD-72260) and the low resolution full-length RCR complex map were derived. **B**) Local resolution maps, Fourier shell coefficient (FSC) curves and electron potential maps of example Fab:RIPR interfaces for RIPR:RP.093:RP.073:RP.063 (PDB ID 9Q6B, EMD-72265) and RIPR:RP.047:RP.057:RP.035 (PDB ID: 9Q69, EMD-72260). **C**) Low resolution map of the RCR:RP.047:RP.057:RP.035:RP.063:RP.073:RP.093 used to build the full-length composite model. **D**) Representative micrograph and 2D classes of the PCRCR:RP.052:RP092 complex sample from which the map and model for RIPR:RP.092:RP.052 (PDB ID 9Q7C, EMD-72294), and the low resolution RIPR:RP.092:RP.052:CSS complex map were derived. **E**) Local resolution map and FSC curves for RIPR:RP.092:RP.052 (PDB ID 9Q7C, EMD-72294). **F**) Low resolution map of the RIPR:RP.092:RP.052:CSS complex used to build the RIPR:CSS composite model.

**Figure S4.**
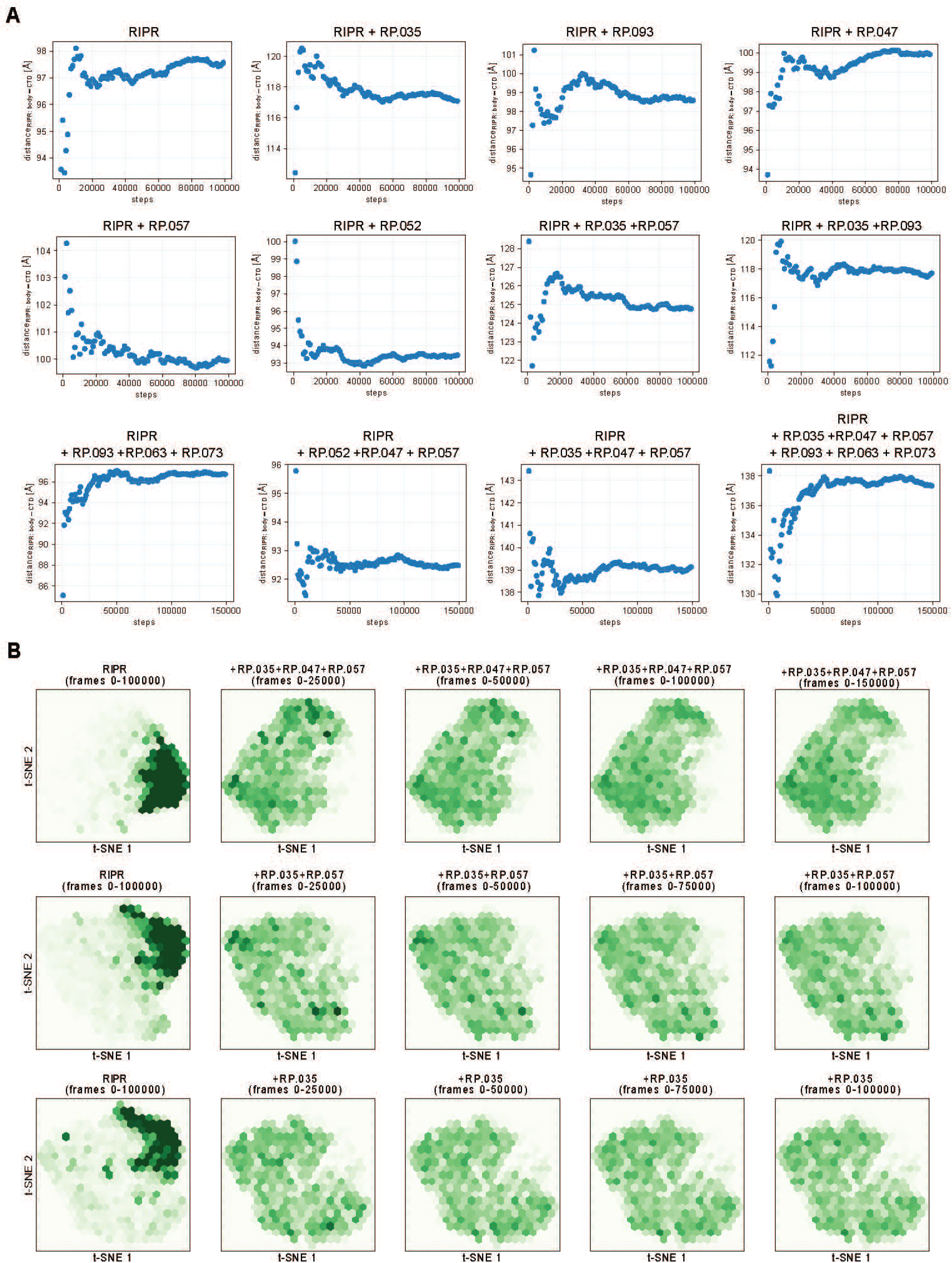
Convergence of the MD trajectories; related to Figure 5. **A**) Cumulative average distance between the centre of mass (COM) of the RIPR-Body and the centre of the target (CDT) is shown as a function of simulation time for each simulated system. The average was recomputed every 1,000 steps. **B**) The conformational space sampled by the RIPR-Tail during the MD simulation is shown in two-dimensional reduced space using t-distributed stochastic neighbour embedding (t-SNE). The panels on the left show the RIPR simulation space for reference. Moving from left to right, different simulation times have been used to evaluate the conformational space explored by the RIPR-Tail.

**Figure S5.**
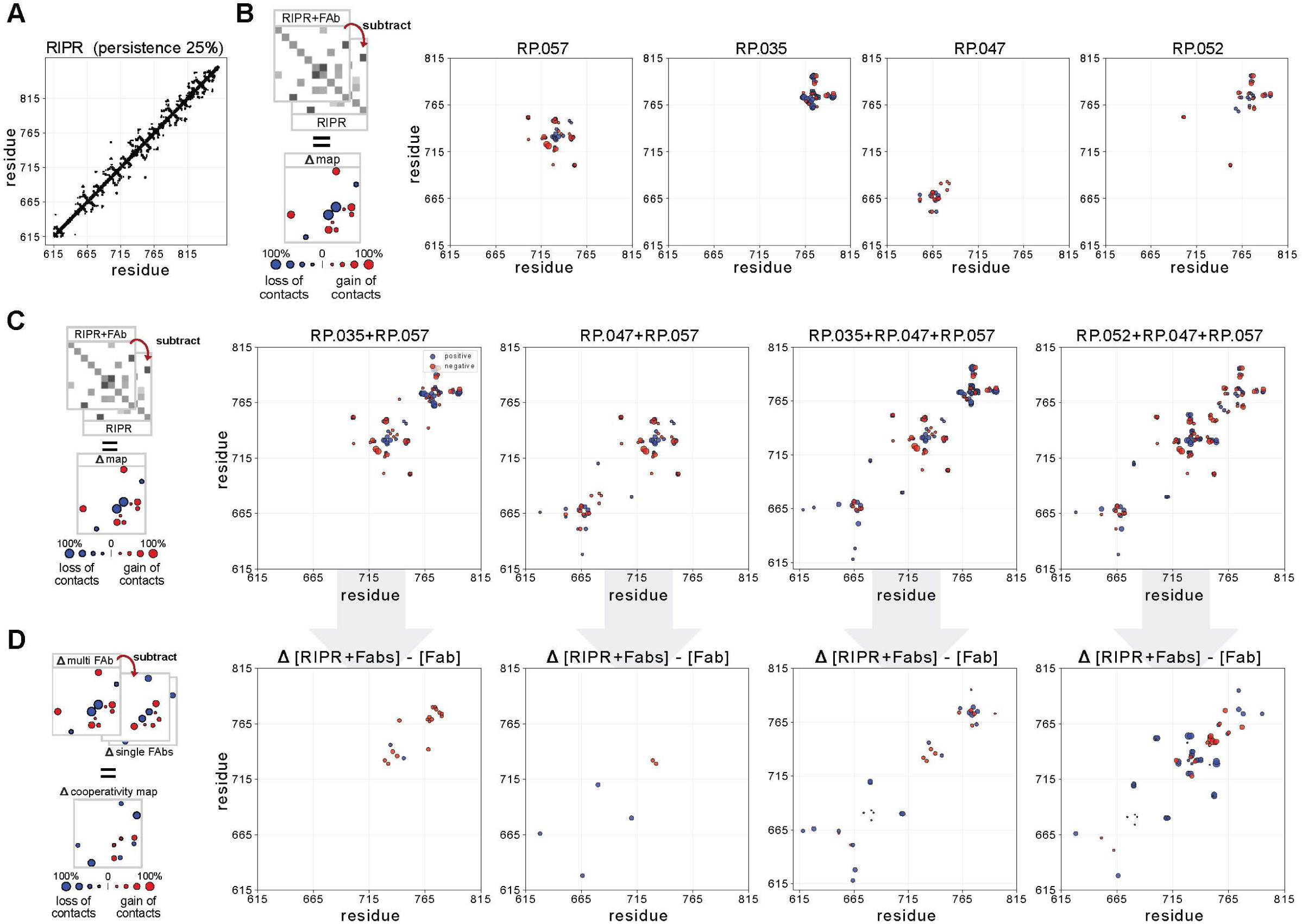
Analysis of the synergistic effects of anti-RIPR^EGF (6-8)^ Fabs on the flexibility of the RIPR-Tail; related to Figure 5. **A**) Contact residue map for simulated RIPR with CALVADOS 3. Only contacts with a persistence of over 25% are shown. **B**) Each plot shows the difference (Δ) in intra-molecular contact maps between simulations of RIPR in a complex with the anti-RIPR^EGF (6-7)^Fabs (RP.047 or RP.057) or the anti-RIPR^EGF (8)^ Fabs (RP.035 or RP.052) versus simulations of RIPR alone. The colour of each point indicates if the contact is lost in the complex (blue) or gained in the complex (red) as compared to the RIPR-alone simulation. The size of each point reports on the magnitude of the contact difference. The procedure for creating the difference contact map is shown schematically on the left of the panel. **C**) Each plot in the panel reports the intra-molecular contacts present in the RIPR simulation with multiple Fabs, and **D**) those that are extra as compared to the RIPR simulation with a single Fabs, to explore possible synergistic effects. The colour of each point indicates if the contact is lost in the complex (blue) or gained in the complex (red) as compared to the RIPR - single Fab simulations. The size of each point reports on the magnitude of the contact difference. The procedure for creating the difference contact map is shown schematically on the left of the panel.

**Figure S6.**
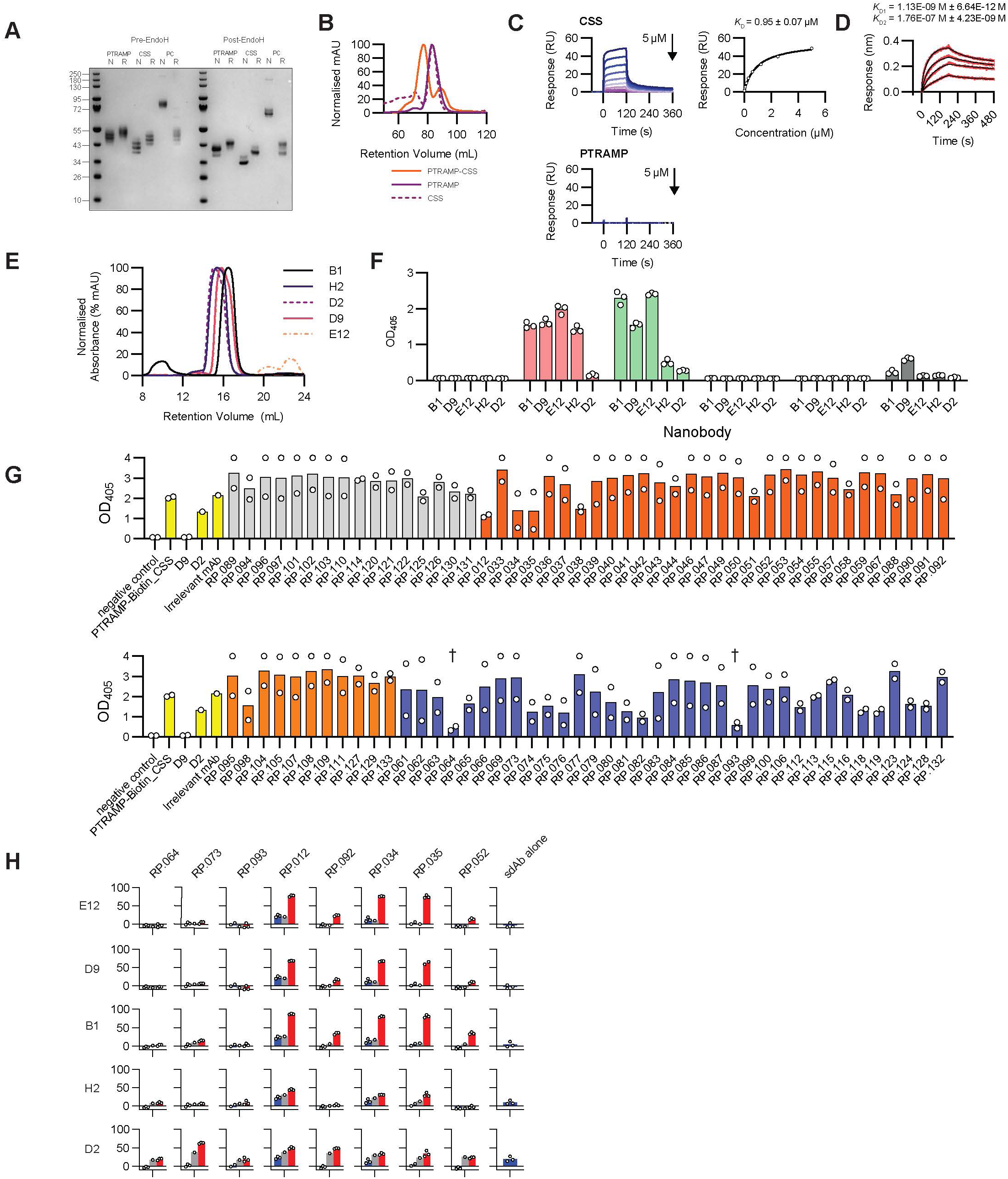
Analysis of the recombinant PTRAMP-CSS-RIPR interaction; related to Figure 6. **A**) Coomassie gel showing PTRAMP, CSS, and PTRAMP-CSS (PC) pre- and post-EndoH treatment under non-reducing (N) and reducing (R) conditions. **B**) Size exclusion chromatograms of PTRAMP, CSS, and PTRAMP-CSS. **C**) SPR sensorgrams show single-cycle kinetics of CSS (left) or PTRAMP (right) binding to mono-biotinylated RIPR-Tail coated CAP sensor ship (blue). Kinetic parameters were determined by Biacore X100 evaluation software by fitting to a bivalent binding model (black). **D**) Representative biolayer interferometry sensorgrams of RIPR binding to PTRAMP-CSS (orange). Data were fitted with a 2:1 heterogenous ligand model (black). **E**) Size exclusion chromatograms of anti-CSS sdAbs. **F**) ELISA results of CSS-sdAbs binding to PTRAMP, CSS, or PTRAMP-CSS heterodimer under normal conditions or following reduction of the target antigens by boiling in Nu-PAGE reducing agent. **G**) ELISA data to assess antibody-mediated blockade of the RIPR : PTRAMP-CSS interaction. Plates were coated with RIPR followed by addition of test anti-RIPR mAb and mono-biotinylated PTRAMP-CSS heterodimer. Bound mono-biotinylated PTRAMP-CSS was detected by ExtrAvidin® -alkaline phosphate. Yellow bars: controls, negative = no mono-biotinylated PTRAMP-CSS, D9 & D2 were pre-incubated with mono-biotinylated PTRAMP-CSS, an unrelated mAb was added at the same time as the RIPR mAbs; grey bars: anti-RIPR Body mAbs; orange bars: anti-RIPR^EGF (5-8)^ mAbs; and blue bars: anti-RIPR^EGF (9-10)-CTD^ mAbs. Negative control was non-biotinylated PTRAMP-CSS. † Indicates mAbs considered blocking. Bars are mean of two independent replicates. **H**) GIA analysis of anti-CSS sdAb and anti-RIPR mAb combinations. Each anti-RIPR mAb was tested alone (1 mg/mL) or in combination with an anti-CSS sdAb (1 mg/mL each). Blue represents the test sdAb or mAb alone, grey the predicted % GIA (Bliss additivity), and red the measured % GIA result.

**Table S1.**
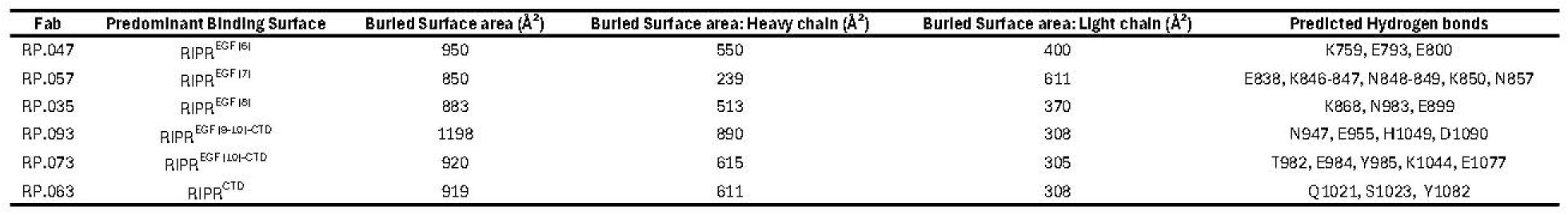
Details of RIPR-Fab binding sites as determine by cryo-EM; related to Figure 3.

| Fab | Predominant Binding Surface | Buried Surface area (Å<sup>2</sup>) | Buried Surface area: Heavy chain (Å<sup>2</sup>) | Buried Surface area: Light chain (Å<sup>2</sup>) | Predicted Hydrogen bonds |
| --- | --- | --- | --- | --- | --- |
| RP.047 | RIPR<sup>EGF (6)</sup> | 950 | 550 | 400 | K759, E793, E800 |
| RP.057 | RIPR<sup>EGF (7)</sup> | 850 | 239 | 611 | E838, K846-847, N848-849, K850, N857 |
| RP.035 | RIPR<sup>EGF (8)</sup> | 883 | 513 | 370 | K868, N983, E899 |
| RP.093 | RIPR<sup>EGF (9-10)-CTD</sup> | 1198 | 890 | 308 | N947, E955, H1049, D1090 |
| RP.073 | RIPR<sup>EGF (10)-CTD</sup> | 920 | 615 | 305 | T982, E984, Y985, K1044, E1077 |
| RP.063 | RIPR<sup>CTD</sup> | 919 | 611 | 308 | Q1021, S1023, Y1082 |

**Data S1 – Antibody germline, epitope, and kinetic data summary, related to Figure 1.**

ELISA epitope data represents a summary of **Data S2.** Kinetic data represent the mean of two repeats. Cells highlighted in purple indicate *k*_off_ values at the limit of the assay (6 x 10^-5^ s^-1^) as antigen did not dissociate from these mAbs significantly during the 15 minute dissociation window. Cells highlighted in yellow indicated mAbs that the residue deviation exceeds >10% of R_max_ (i.e. curves do not follow 1:1 binding). Cells highlighted in turquoise are due to Max datapoint falling <50% below predicted R_max_ value.

**Data S2 – Details of mAb epitope binding as determined by ELISA, related to Figure 1.**

RIPR protein truncations are ordered left-to-right reflecting N- to C-terminal domains. mAb epitope was determined by the last truncation which yielded an OD comparable to FL-RIPR. Values are the mean of 3 independent repeats.

**Data S3 – Cryo-EM data collection, refinement and validation statistics, related to Figures 3, 4, and 6.**

**Data S4 – Glycoproteomic summary data, related to Figure 6.**

Peptide coverage and occupancy data shown.

## STAR METHODS

### CONTACT FOR REAGENT AND RESOURCE SHARING

Further information and requests for resources should be directed to and will be fulfilled by the Lead Contact, Simon J. Draper.

## Data and Code Availability

Requests for monoclonal antibodies (mAbs) generated in the study should be directed to the Lead Contact, Simon J. Draper. Custom code to reproduce the MD results and figures of this study will be made publicly available before the final version of this manuscript is published. In the meantime, it is available upon request by email to the authors.

## EXPERIMENTAL MODEL AND SUBJECT DETAILS

### Cell Lines

Expi293F HEK cells were cultured in suspension in Expi293 expression medium (Thermo Fisher Scientific) at 37 °C, 8% CO_2_, on an orbital shaker set at 125 RPM. *Drosophila* S2 cells were cultured in suspension in EX-CELL 420 medium (Sigma-Aldrich) supplemented with 100 U/mL penicillin, 0.1 mg/mL streptomycin and 10% foetal bovine serum (FBS) at 25 °C. Stable S2 cell lines expressing RH5 or RIPR proteins were generated using the ExpreS2 platform (ExpreS2ion Biotechnologies, Demark) as previously described.^8,48^

### Experimental Animal Models

Mice were housed under United Kingdom Home Office Project Licence 70/8718 with all procedures carried out receiving approval of the Wellcome Trust Sanger Institute Animal Welfare and Ethical Review Body.

### Human Blood Sample Collection

Human polyclonal total IgG was purified from serum obtained from healthy UK adults vaccinated with R78C/Matrix-M® in the VAC089 Phase 1a clinical trial (NCT05385471; Salkeld J *et al.*, in preparation). The VAC089 trial is a non-randomised, open-label, first-in-human, Phase 1a trial of the blood-stage malaria candidate vaccine R78C, alone or in combination with RH5.1, formulated with Novavax’s Matrix-M® adjuvant. The trial was conducted at the Centre for Clinical Vaccinology and Tropical Medicine at the University of Oxford. The study received ethical approval from the UK NHS Research Ethics Service (London - Harrow Research Ethics Committee, Ref 22/LO/0715) and was approved by the UK Medicines and Healthcare products Regulatory Agency (Ref 21584/0461/001-0001).

Participants signed written consent forms and consent was verified before each vaccination. The trial was registered on ClinicalTrials.gov (NCT05385471) and was conducted according to the principles of the current revision of the Declaration of Helsinki 2008 and in full conformity with the ICH guidelines for Good Clinical Practice (GCP).

## METHOD DETAILS

### Recombinant protein expression and purification

Recombinant full-length RH5, CyRPA and RIPR proteins were expressed as previously reported.^17^ RIPR protein truncations were based on the full-length RIPR sequence with 12 mutations to remove N-linked glycosylation sequons: N103Q, N114Q, N228Q, N334Q, N480Q, N498Q, N506Q, N526Q, N646Q, N647Q, N964Q, N1021Q. The protein truncation boundaries are as follows; RIPR-Tail: D717 – N1086, EGF 6+: K771 – N1086, EGF 7+: D817 – N1086, EGF 8+: E856 – N1086, EGF 9+: L898 – N1086, EGF 10+: Q940 – N1086, CTD: R980 – N1086. Each protein construct included an N-terminal BiP secretion signal and a C-terminal four-amino acid purification tag (C-tag: EPEA^8^). Each RIPR-Tail truncation was expressed as secreted protein by stable polyclonal *Drosophila* S2 cell lines (ExpreS2ion Biotechnologies) as previously reported.^8,48^ All supernatants were harvested via centrifugation and the proteins purified using CaptureSelect C-tag affinity matrix (Thermo Fisher Scientific) on an ÄKTA Pure FPLC system (Cytiva). A further size exclusion chromatography (SEC) polishing step was performed on a HiLoad 16/60 Superdex 200 pg column (GE Healthcare) in 20 mM Tris, 150 mM NaCl, pH 7.4.

Biotinylated RIPR and RIPR-Tail were produced in S2 cell lines as above but including a C-terminal biotin acceptor peptide (BAP) tag (FL-RIPR) or N-terminal BAP-tag (RIPR-Tail). Cell lines were co-transfected with BirA enzyme to produce mono-biotinylated protein.

Full length CSS comprising residues Q21-K290, and full-length PTRAMP comprising residues C30-T307, both with a mIgG secretion signal sequence and C-terminal His6-tag, were expressed using Expi293F HEK cells according to the manufacturer’s instructions. All supernatants were harvested via centrifugation and the proteins purified using HisTrap Excel resin (Cytiva) on an ÄKTA Pure FPLC system (Cytiva). A further SEC polishing step was performed on a HiLoad 16/60 Superdex 200 pg column (GE Healthcare) in 20 mM Tris, 150 mM NaCl, pH 7.4. Post size exclusion, the proteins were treated with Endo H_f_ (NEB) for 2 h at 37 °C. Endo H_f_ was removed using a MBPTrap™ HP column (Cytiva) on an ÄKTA Pure FPLC system (Cytiva). The flow-through containing PTRAMP/CSS was collected and concentrated using an Amicon ultra centrifugal concentrator (Millipore) with a molecular weight cut-off of 10 kDa. PTRAMP-CSS heterodimer was produced by co-transfecting Expi293 cells with PTRAMP and CSS at a 3:1 ratio. All other purification steps remained the same.

### Mouse immunisations

The mouse study was carried out at the Babraham Institute Biological Support Unit under Project License 70/8718 issued by the UK Government Home Office under the Animal (Scientific Procedures) Act (A(SP)A), 1986, incorporating Directive 2010/63/EU of the European Parliament, and with the approval of the Sanger Institute Animal Welfare and Ethical Review Body.

The Kymouse platform mice have a genetic background of a mixture of 129S7 and C57BL/6J strains and contain the entire human immunoglobulin variable-gene repertoire.^31^ All mice were housed on a 12 h light/dark cycle at 20–24 °C with 55% ± 10% humidity. Groups of 6 male and female 7–12-week-old Kymouse platform mice were immunised with either 10 μg recombinant RIPIR protein or 8 μg preformed RCR-complex (recombinant RH5, CyRPA and RIPR proteins) formulated 1:1 vol/vol with AddaVax adjuvant (InvivoGen). Mice were injected intramuscularly into the tibialis muscle of each hind leg using a 30-gauge needle (BD) with 25 μL per injection site (50 μL total) of immunogen under isoflurane anaesthesia. Each mouse received a dose on days 0, 14, 42 and 56. Mice were sacrificed on day 61 under UK Home Office Schedule 1 (rising concentration of CO_2_) and spleens were processed to generate single-cell suspensions that were cryopreserved in 10% DMSO in heat-inactivated FBS and stored in liquid nitrogen until further analysis. Pre-bleed, serial bleeds and terminal bleeds were collected for serological analysis. Whole blood (0.02 mL) was collected via tail vein on days -7, 7, 35 and 49. Terminal bleeds (0.1 mL) were collected at the time of sacrifice via cardiac bleed. Serum was separated from haematocrit via centrifugation at 2000 x g for 10 min. Serum was stored at −20 °C and was used to monitor polyclonal antibody titres by ELISA.

### B cell isolation and sequencing from Kymouse Platform mice

Splenocytes were thawed and centrifuged at 400 × g for 10 min at 4 °C and resuspended in FACS buffer (PBS, 1% FBS, 1 mM EDTA, 25 mM HEPES). Splenocytes were stained for B220 (BUV395), CD19 (BUV395), IgM (BV605), IgG (BUV737), IgD (BV510), and for Ly-6G, CD8α, F4/80 and CD4 (all APC-Cy7), as well as either FL-RIPR-biotin :: streptavidin-AF488 or RIPR^CTD^-biotin :: streptavidin-PE and RIPR^EGF(5-8)^-biotin :: streptavidin AF488 antigens in Brilliant buffer (Invitrogen). Single B220/CD19+ IgM- IgD- IgG+ AF488+ PE+ B cells were sorted by FACS using a BD FACS Aria Fusion flow cytometer (Beckton Dickinson) and collected into individual wells of 96-well plates containing QuickExtract RNA Extraction lysis buffer (Epicenter).

Variable heavy (VH) and variable light (VL) paired sequences of Ig genes were amplified by reverse-transcription (WarmStart RTx Reverse Transcriptase, NEB) followed by PCR (Q5 DNA polymerase, NEB). Ilumina sequencing libraries were generated and sequenced using an Ilumina MiSeq (MiSeq Reagent Kit v3, Illumina). Antibody sequence data were analysed using IntelliSellect software and clones were selected from the sequencing data combined with the FACS index data.

### Antibody expression and purification

Antibody heavy and light chains were ordered and cloned into AbVec expression plasmids to produce recombinant human IgG1 mAbs^49^ (a gift from Patrick C. Wilson, University of Chicago, USA) by Twist Biosciences, USA. Anti-RIPR mAbs were transiently expressed in Expi293F HEK cells. Cognate heavy and light chain-coding plasmids were co-transfected at a 1:1 ratio. Supernatants were harvested via centrifugation. All mAbs were purified using a 5 mL Protein A HP column (Cytiva) on an ÄKTA Pure FPLC system (Cytiva). Equilibration and wash steps were performed with PBS supplemented with 500 mM NaCl, mAbs were eluted in 0.1 M glycine pH 2.7, 100 mM NaCl.^50^ The eluates were pH equilibrated to 7.4 using 1.0 M Tris HCl pH 9.0 and immediately buffer exchanged into Dulbecco’s PBS (DPBS) and concentrated using an Amicon ultra centrifugal concentrator (Millipore) with a molecular weight cut-off of 30 kDa. Total IgG from mouse serum was purified using the same method. For Fabs, the heavy chain was modified to terminate at the end of the CH1 domain, and expressed in the same manner as mAbs. Purification was performed using CaptureSelect™ CH1-XL Affinity Matrix (Thermo Fisher Scientific) in gravity flow columns. Equilibration and wash steps were performed with Tris-buffered saline (TBS) and Fabs were eluted with 50 mM sodium acetate, pH 4.0 and immediately buffer exchanged into TBS and concentrated using an Amicon ultra centrifugal concentrator (Millipore) with a molecular weight cut-off of 10 kDa.

### Human polyclonal IgG purification

Serum samples from 28 days post**-**final vaccination from participants vaccinated with three doses of 10 µg R78C protein formulated with 50 μg Matrix-M adjuvant (ClinicalTrials.gov NCT05385471) were pooled and purified using a 5 mL Protein A HP column (Cytiva) on an ÄKTA Pure FPLC system (Cytiva) as described above.

### Anti-RIPR^EGF (5-8)^-specific IgG depletion from rabbit anti-RIPR IgG

Recombinant RIPR^EGF (5-8)^ protein^17^ was coupled to a HiTrap NHS-Activated HP affinity column (Cytiva) using standard amine coupling protocols. Anti-full length RIPR polyclonal total IgG from rabbits generated in a previous study^17^ was run over the column using an ÄKTA Pure FPLC system (Cytiva) and the flow through collected as the RIPR^EGF (5-8)^-depleted fraction.

RIPR^EGF (5-8)^-specific IgG was eluted in 0.1 M glycine, pH 2.7, followed by pH equilibration to 7.4 using 1.0 M Tris HCl, pH 9.0, and immediately buffer-exchanged into incomplete parasite growth media (RPMI, 2 mM L-glutamine, 0.05 g/L hypoxanthine, 5.94 g/L HEPES) and concentrated using an Amicon ultra centrifugal concentrator (Millipore) with a molecular weight cut-off of 30 kDa.

### Assay of growth inhibition activity (GIA)

GIA assays against 3D7 clone *P. falciparum* parasites were carried out over one blood-stage growth cycle (∼48 h) as previously described.^51^ Briefly, the assay was performed at indicated concentrations of mAb or purified total IgG in triplicate wells and a biochemical measurement using *P. falciparum* lactate dehydrogenase assay was used to quantify parasitaemia and define % growth inhibition. To ensure consistency between experiments, in each case the activity of a negative control mAb, EBL040^52^, which binds to the Ebola virus glycoprotein, and three anti-RH5 mAbs with well-characterised levels of GIA (2AC7^16^, R5.016^22^, R5.034^21^) were run alongside the test samples and used for assay QC. Purified total IgG samples from immunised animals were pre-incubated with human group O RhD-positive red blood cells to eliminate spurious GIA results caused by hemagglutination. Test antibodies were buffer exchanged into incomplete parasite growth media (RPMI, 2 mM L-glutamine, 0.05 g/L hypoxanthine, 5.94 g/L HEPES, 2g/L d-glucose) before performing the GIA assay. To assess for synergistic GIA, mAbs or purified total IgGs were assessed by measuring the GIA activity of an equal mix of two or more antibodies. The predicted Bliss additivity GIA was determined based on the measured activity from each antibody alone using formulas previously described.^32^ For GIA-reversals, the experiment was carried out as above using a fixed amount of antigen-specific IgG. In addition, purified full-length proteins or protein fragments were titrated into the assay with the IgG at a defined concentration, typically between 4 and 5 µM. Each protein was tested in the absence of test IgG as a control.

Red-blood cells for ongoing *Pf*3D7 culture was obtained through either in-house donation or via NHSBT. All human blood donations and purchases at the University of Oxford for use in the GIA assay are anonymised and covered under ethical approval from the Health Research Authority (REC reference 18/LO/0415, protocol number OVC002).

### ELISA

For the assessment of ELISA titres in **Figure S1B,C**, Maxisorp plates were coated overnight at 4 °C (>16 h) with either 2 µg/mL full-length RIPR or an equimolar amount totalling 2 µg/mL of RH5, CyRPA and RIPR (RCR-complex) which had been pre-incubated for 20 min at room temperature (RT). Plates were washed in wash buffer (PBS with 0.05% Tween 20 [PBST]) and blocked with 200 µL/well of Blocker™ Casein (Thermo Fisher Scientific) for 1 h. Plates were washed and antibodies at 10 µg/mL diluted in casein were added. Following a 2 h incubation, plates were washed and a 1:2000 dilution of goat anti-human IgG (γ-chain specific) alkaline phosphate conjugate antibody (A3187, Thermo Fisher) was added and incubated for 1 h.

Plates were washed in washing buffer and 100 µL development buffer added (p-nitrophenyl phosphate substrate diluted in diethanolamine buffer) and developed according to internal controls. A standard curve and Gen5 ELISA software v3.04 (BioTek, UK) were used to convert the optical density 405 nm of individual test samples into arbitrary units (AU).

For assessment of anti-RIPR mAb antibody binding by ELISA in **Data S2**, Nunc Maxisorp plates were coated overnight at 4 °C (>16 h) with either full-length RIPR or a RIPR truncation as indicated at 2 µg/mL. Plates were washed in wash buffer (PBST) and blocked with 200 µL/well of Blocker™ Casein (Thermo Fisher Scientific) for 1 h. Plates were washed and antibodies at 2 µg/mL diluted in casein were added. Following a 1 h incubation, plates were washed and a 1:2000 dilution of goat anti-human IgG (γ-chain specific) alkaline phosphate conjugate antibody (A3187, Thermo Fisher) was added and incubated for 1 h. Plates were washed in washing buffer and 100 µL development buffer added (p-nitrophenyl phosphate substrate diluted in diethanolamine buffer) and optical density read at 405 nm on an Infinite 50 plate reader (Tecan). Unless otherwise stated 50 µL was added per well and all steps were carried out at RT.

For sdAb binding ELISA (**Figure S5F**), 96-well Nunc Maxisorp plates were coated overnight at 4 °C (>16 h) with 2 μg/mL PTRAMP, CSS, or PTRAMP-CSS in both reduced, by heating to 90°C for 10 minutes in Nu-Page 10x reducing buffer, and non-reduced forms. Plates were washed in wash buffer (PBST) and blocked with 200 µL/well of Starting Block™ T20 (Thermo Fisher Scientific) for 1 h. Plates were washed, and sdAbs at 2 µg/mL diluted in PBS were added.

Plates were washed then incubated for 1 h with 50 μL/well MonoRab™ Rabbit Anti-Camelid VHH Cocktail [Biotin] at 2μg/mL. Plates were washed, and a 1:2000 dilution of ExtrAvidin®–Alkaline Phosphatase (E2636, Merck) was added and incubated for 1 h. Plates were washed and 100 µL development buffer added (p-nitrophenyl phosphate substrate diluted in diethanolamine buffer) and optical density read at 405 nm on an Infinite 50 plate reader (Tecan). Unless otherwise stated 50 µL was added per well and all steps were carried out at RT.

For assessment of RIPR-PTRAMP-CSS blocking (**Figure S5G**), Nunc Maxisorp plates were coated overnight at 4 °C (>16 h) with full-length RIPR at 2 µg/mL. Plates were washed in wash buffer (PBST) and blocked with 200 µL/well of Blocker™ Casein (Thermo Fisher Scientific) for 1 h. Plates were washed and antibodies at 2 µg/mL diluted in casein were added. Following a 1 h incubation, plates were washed and 2 µg/mL mono-biotinylated PTRAMP-CSS was added and incubated for 1 h. As controls, mono-biotinylated PTRAMP-CSS pre-incubated with 2 µg/mL sdAbs was added. Plates were washed, and a 1:4000 dilution of ExtrAvidin®–Alkaline Phosphatase (E2636, Merck) was added and incubated for 1 h. Plates were washed and 100 µL development buffer added (p-nitrophenyl phosphate substrate diluted in diethanolamine buffer) and optical density read at 405 nm on an Infinite 50 plate reader (Tecan). Unless otherwise stated 50 µL was added per well and all steps were carried out at RT. For each ELISA a positive control was included in each plate (biotinylated PTRAMP-CSS with no mAb/sdAb bound), this was developed to an OD_405_ of 2. A negative control was also included in each assay where a non-biotinylated analyte was added with no mAb/sdAb bound.

For assessment of RIPR-PTRAMP-CSS dissociation (**Figure 6F**): Nunc Maxisorp plates were coated overnight at 4 °C (>16 h) with full-length RIPR at 2 µg/mL. Plates were washed in wash buffer (PBST) and blocked with 200 µL/well of Blocker™ Casein (Thermo Fisher Scientific) for 1 h. Plates were then coated with 2 µg/mL mono-biotinylated PTRAMP-CSS for 1 h. Plates were washed and then antibody mixes were added using a 5-fold dilution series starting at 50 µg/mL (16.6 µg/mL of each mAb) diluted in casein. Following a 1 h incubation, plates were washed and a 1:4000 dilution of ExtrAvidin®–Alkaline Phosphatase (E2636, Merck) was added and incubated for 1 h. Plates were washed and 100 µL development buffer added (p-nitrophenyl phosphate substrate diluted in diethanolamine buffer) and optical density read at 405 nm on an Infinite 50 plate reader (Tecan). Unless otherwise stated 50 µL was added per well and all steps were carried out at RT.

### High throughput antibody kinetics

High-throughput SPR binding experiments (**Data S1**) were performed on a Carterra LSA instrument equipped with a 2D planar carboxymethyldextran surface (CMDP) chip type (Carterra) using a 384-ligand array format. The CMDP chip was first conditioned with 60 s injections of 50 mM NaOH, 1 M NaCl and 10 mM glycine (pH 2.0) before activation with a freshly prepared 1:1:1 mixture of 100 mM MES (pH 5.5), 100 mM sulfo-N-hydroxysuccinimide, and 400 mM 1-ethyl-3-(3-dimethylaminopropyl) carbodiimide hydrochloride. A coupled lawn of goat anti-human IgG Fc (hFc; 50 µg/mL in 10 mM sodium acetate, pH 4.5) (Jackson ImmunoResearch) was then prepared before quenching with 0.5 M ethanolamine (pH 8.5) and washing with 10 mM glycine (pH 2.0). mAbs prepared at 1 µg/mL in Tris-buffered saline with 0.01% Tween-20 (TBST) were then captured onto the surface in a 384-array format via a multi-channel device, capturing 96 ligands at a time. For binding kinetics and affinity measurements, a six-point three-fold dilution series of RIPR protein (33.3 nM – 46.7 pM) in TBST was sequentially injected onto the chip from lowest to highest concentration. For each concentration, the antigen was injected for 5 min (association phase), followed by TBST injection for 15 min (dissociation phase). Two regeneration cycles of 30 s were performed between each dilution series by injecting 10 mM glycine (pH 2.0) on the chip surface. The SPR results were exported to Kinetics Software (Carterra) and analysed as non-regenerative kinetics data to calculate association rate constant (*k*_on_), dissociation rate constant (*k*_off_) and equilibrium dissociation constant (*K*_D_) values via fitting to the Langmuir 1:1 model. Prior to fitting, the data were referenced to the anti-hFc surface then double referenced using the final stabilising blank injection. Any mAb that did not conform to the Langmuir 1:1 model (defined as having a residual standard deviation >7% of the calculated R_max_) was termed ‘Non-1:1’ and is highlighted in yellow in **Data S1**; this comprised 11/92 of the mAbs tested.

### Epitope binning

Epitope binning experiments shown in **Figure 1D, S1F,G** were performed by high-throughput surface plasmon resonance (SPR) in a classical sandwich assay format using the Carterra LSA and an HC200M chip. The chip was conditioned as described above before antibodies prepared at 10 µg/mL in 10 mM sodium acetate (pH 4.5) with 0.05% Tween were coupled to the surface: the chip surface was first activated with a freshly prepared 1:1:1 activation mix of 100 mM MES (pH 5.5), 100 mM sulfo-N-hydroxysuccinimide, and 400 mM 1-ethyl-3-(3-dimethylaminopropyl) carbodiimide hydrochloride, and antibodies were injected and immobilised onto the chip surface by direct coupling. The chip surface was then quenched with 1 M ethanolamine (pH 8.5), followed by washing with 10 mM glycine (pH 2.0). Sequential injections of 100 nM RIPR protein (5 min) followed immediately by the 10 µg/mL sandwiching antibody (5 min), both diluted in HEPES-buffered saline Tween-EDTA (HBSTE) with 0.5 mg/mL bovine serum albumin (BSA), were added to the coupled array and the surfaces regenerated with either 10 mM NaOH (pH 10) using two 30 s regeneration cycles for the anti-RIPR^EGF (5-8)^ mAbs, and 10 mM glycine pH 2.1 + 0.8 M MgCl_2_ using two 30 s regeneration cycles for the anti-RIPR^EGF (9-10)-CTD^ mAbs. Data were analysed using the Carterra Epitope software. Two separate experiments were run to reduce the number of antibodies analysed in each run thus reducing the number of regeneration cycles.

### Surface plasmon resonance

SPR was carried out using the Biacore™ X100 machine and software. RIPR, RIPR-Tail truncations, or PTRAMP-CSS were immobilised onto separate CM5 Sensor Chips (Cytiva) using the standard amine coupling protocol, yielding ∼900 response units (RU) for each antigen. RIPR, RIPR-Tail truncations, PTRAMP-CSS, PTRAMP, or CSS were diluted in SPR buffer (PBS+P20: 137 mM NaCl, 2.7 mM KCl, 10 mM Na_2_HPO_4_, 1.8 mM KH_2_PO_4_, 0.005 % surfactant P20 (Cytiva)) to yield a top concentration of 5 or 50 µM. For each protein, a two-fold serial dilution series was then prepared in the same buffer. Samples were injected for 180 s at 30 μL/min before dissociation for 700 s. The chip was regenerated with a 30 s injection of 10 mM glycine pH 1.5. Data were analysed using the Biacore X100 Evaluation software v2.0.2, and the equilibrium dissociation constant (*K*_D_) was determined from a plot of steady-state binding levels.

### Biolayer interferometry

Biolayer interferometry (BLI) experiments (**Figure S6D**) were carried out on an Octet® RED384 System (ForteBio) at 25 °C. Octet® AHC Biosensors (Sartorius) were used. Biosensors were initially dipped in kinetics buffer for 180 s to establish a baseline signal, and then dipped into wells containing mAb RP.103 diluted to 20 µg/mL kinetic buffer (TBS + 0.05% Tween), followed by another 180 s baseline in kinetic buffer. The biosensors were then dipped into wells containing RIPR diluted to 25 µg/mL for 300 s. This was followed by a further dip in kinetic buffer for 180 s. The biosensors were then dipped in PTRAMP-CSS diluted to 200 nM in kinetic buffer, or a kinetic buffer reference, and association measured for 180 s. Dissociation was measured by dipping into kinetics buffer for 420 s. Data were analysed using Sartorius Data Analysis software HT 11.1.1.39. Kinetic curves were fitted using a 2:1 heterogenous ligand model.

### Mass photometry

Mass photometry (MP) measurements were conducted using a Refeyn TwoMP system as previously described.^53^ High Precision No. 1.5H glass coverslips were cleaned via sonication in Milli-Q H_2_O, followed by isopropanol and Milli-Q H_2_O then dried under nitrogen flow. Sample chambers were assembled using silicone gaskets (CultureWell™ reusable gasket, 3 mm diameter x 1 mm depth, Grace Bio-Labs). Coverslips were placed on the sample stage and a single gasket was filled with 5-20 µL DPBS (without calcium, without magnesium, pH 7.4 Thermo Fisher Scientific) to find focus and ensure low background signal-to-noise. Samples were measured at a final concentration between 10 and 50 nM.

Acquisition settings within AcquireMP (v2023 R1.1, Refeyn Ltd) were as follows: regular field of view, frame binning = 2, frame rate = 498.3 Hz, pixel binning = 6, exposure time 1.95 ms and movies were taken over 60 s. Mass calibration was conducted using an in-house protein standard. Data were analysed using DiscoverMP (v2024 R2.1, Refeyn Ltd). Molecule counts were used to determine levels of protein complexes. Representative histograms were generated using DiscoverMP.

### Glycoproteomics

Approximately 5 µg protein was loaded and run on an SDS-PAGE. Gel bands were excised and washed sequentially with HPLC grade water followed by 1:1 (v/v) MeCN/H_2_O. Gel bands were dried (via vacuum centrifuge), treated with 10 mM dithiothreitol (DTT) in 100 mM NH_4_HCO_3_ and incubated for 45 min at 56 °C with shaking. DTT was removed and 55 mM iodoacetamide (in 100 mM NH_4_HCO_3_) was added and incubated for 30 min in the dark. All liquid was removed and gels were washed with 100 mM NH_4_HCO_3_/MeCN as above. Gels were dried and 12.5 ng/µL trypsin, chymotrypsin or alpha lytic protease was added separately and incubated overnight at 37 °C. Samples were then washed and (glyco)peptides were extracted and pooled with sequential washes with 5% (v/v) formic acid (FA) in H_2_O and MeCN. Dried samples were reconstituted in 0.05% trifluoroacetic acid and run by LC-MS.

Samples were analysed using an Ultimate 3000 UHPLC coupled to an Orbitrap Q Exactive mass spectrometer (Thermo Fisher Scientific). Peptides were loaded onto a 75 µm × 2 cm pre-column and separated on a 75 µm × 15 cm Pepmap C18 analytical column (Thermo Fisher Scientific). Buffer A was 0.1% FA in H_2_O and buffer B was 0.1% FA in 80% MeCN with 20% H_2_O. A 120 min linear gradient (0% to 40% buffer B) was used. A universal HCD identification method was used. Data were collected in data-dependent acquisition mode with a mass range 375 to 1500 m/z and at a resolution of 70,000. For MS/MS scans, stepped HCD normalised energy was set to 27, 30 and 33% with orbitrap detection at a resolution of 35,000.

Data were analysed in Byos (Protein Metrics). Digestion was set to RK, FYWML and TASV for trypsin, chymotrypsin and alpha-lytic protease digests, respectively, with maximum of two missed cleavages. Fixed modifications were carbamidomethylation (57.02 Da) and variable modifications were methionine oxidation (15.99 Da). Byos in-built common human N-linked (59 glycans) glycan library was used. FDR was set to 0.01 and mass tolerances for precursor and fragment ions set to 6.00 ppm and 20.00 ppm respectively. For analysis, a minimum Byos threshold score of 400 was used for glycopeptide filtering, with the exception of peptides covering site PTRAMP N226 for which a score of 150 was used. Glycopeptides were manually validated. For quantification, the extracted ion chromatogram intensities for each glycopeptide of that sequence were summed and plotted relative to the total intensity for each glycosite.

### SdAb production

SdAb sequences^14^ were subcloned into pET15b vectors containing a HIS-tag for expression and transformed into BL21 (DE3) cells for *Escherichia coli* expression. 500 mL Luria-Bertani (LB) was inoculated with 10 mL starter culture, supplemented with 0.8% w/v glucose and grown at 37 °C, 200 rpm until the OD_600_ reached 0.6-0.8. Protein expression was then induced with IPTG (0.42 mM), cultures incubated for 4 h at 30 °C, 200 rpm. Cell pellets were harvested via centrifugation for 15 min at 9000 x g, 25 °C. The sdAbs were released from the pelleted cells using osmotic lysis. Cell pellets were resuspended with 15 mL ice-cold TES buffer (0.2 M Tris-HCL, 0.51 mM EDTA, 0.5 M sucrose, pH 8.0) and incubated for 1 h on a tube shaker at 4 °C. Cells were then centrifuged for 15 min at 9000 x g, 25 °C, and the supernatant was retained. 30 mL TES/4 (0.05 M Tris-HCL, 0.13 mM EDTA, 0.125 M sucrose, pH 8.0) was added, incubated for 45 min on a tube shaker at 4 °C. The samples were then centrifuged for 30 min at 10,000 x g, 4 °C and the supernatant retained. All sdAbs were purified using HisTrap™ HP columns (Cytiva) on an ÄKTA Pure FPLC system (Cytiva). A further SEC polishing step was performed on a HiLoad 16/60 Superdex 200 pg column (GE Healthcare) in 20 mM Tris, 150 mM NaCl, pH 7.4.

### Cryo-EM sample preparation and data collection

Individual PCRCR-complex components were mixed at a 1:1 molar ratio at a final concentration of 1.31 mg/mL in TBS and incubated for 10 min at RT. Fabs were added to the PCRCR-complex mixture at a 1:1 ratio at a final total protein concentration of 0.8 mg/mL in TBS and incubated for 10 min at RT. RvLEAMshort was produced as published and added at a final concentration of 0.625 mg/mL at the beginning of the incubation to improve protein orientation distribution.^54,55^ 3 μL each complex was added to UltrAuFoil R 1.2/1.3 300 mesh gold grids which had been plasma washed by glow discharging with Leica Coater ACE200 for 30 s at 10 mA. The protein was plunge frozen in liquid ethane using Vitrobot Mark IV (Thermo Fisher) after incubation for 5s, blotted for 5 s at 4 °C, 100% humidity. The grids were then stored in liquid nitrogen for data collection.

Grids were loaded into a Glacios 2 microscope (Thermo Fisher Scientific) operating at 200 kV with a Falcon 4i direct electron detector, except for the PCRCR:RP.092:RP.052 dataset, which was collected with a Glacios microscope (Thermo Fisher Scientific) operating at 200 kV with an FEI Falcon IV direct electron detector. Automated data collection for all complexes was carried out using EPU (Thermo Fisher Scientific) at a nominal magnification of 190,000, with a total exposure dose of ∼45 e-/Å2 at a pixel size 0.718 Å (0.725 Å for the PCRCR:RP.092:RP.052 dataset), with a nominal defocus range of -0.8 to -2 μm. Full data collection parameters for each map and model are show in **Data S3**.

### Cryo-EM data processing and model building

All datasets were processed using cryoSPARC^56^. Dose-weighted movie frame alignment was carried out using Patch motion correction in cryoSPARC live to account for stage drift and beam-induced motion. The contrast transfer function (CTF) was estimated using Patch CTF in cryoSPARC live. Micrographs with a CTF fit >10 Å were excluded. Patch motion correction was performed on a subset of 100 micrographs to generate data for micrograph denoising. The micrograph denoise job was performed on these 100 micrographs, and the resulting denoise model was used to denoise the entire micrograph stack. Particles were selected from denoised micrographs using blob picker and extracted. 2D classification was performed and particles from good classes were used for ab-initio reconstruction, followed by heterogenous refinement. Particles from the best heterogenous refinement class for each region of the complex were selected and subjected to 3D classification. Particles from the highest resolution and least orientation biased 3D classes were selected and the particles and best 3D volume were used for a subsequent non-uniform refinement job.^54^ AlphaFold3-predicted models of the antibody Fv regions and the RIPR protein were fitted into the non-uniform refinement volume in UCSF ChimeraX^57^. Regions of the RIPR protein for which no density was seen were deleted, to generate an initial model. The initial model was used to generate a 15 Å molmap, which was imported into cryoSPARC and used for mask generation. Global and local CTF refinement was performed on the final particle stack, followed by local refinement using the generated mask, to generate the final map. The models were manually adjusted using Coot^58^ and further refined through real-space refinement in Phenix^59^ using the final local-refinement map output. The antibody Fv chains were renumbered based on the Kabat numbering scheme in Phenix. Buried surface area (BSA), and hydrogen bonds were analysed using the PISA server65 and ChimeraX. BSA was calculated using a cutoff of >5 Å^2^, and hydrogen bonds were defined as atomic interactions with a distance of ≤3.5 Å. For generation of the composite model of the full RCR-complex, the final high-resolution models of RIPR:RP.047:RP.057:RP.035 (PDB ID: 9Q69, EMD-72260), RIPR:RP.093:RP.073:RP.063 (PDB ID 9Q6B, EMD-72265) and the published model of the RH5-CyRPA-RIPR^Body^ complex (PDB ID: 8CDD) were fitted into a map of the entire complex in UCSF ChimeraX. Structural figures were generated using UCSF ChimeraX.^57^

### Molecular dynamics (MD) simulation

The RIPR and Fab structures used for the MD simulations were extracted from the cryo-EM RCR:RP.047:RP.057:RP.035:RP.093:RP.073:RP.063 complex structure. For the RP.052 simulations, the structure was modelled in ChimeraX by extracting the RP.052 structure and its relative position to RIPR from the RCR:RP.092:RP.052:CSS complex cryo-EM structure and then fitted to the RCR:RP.047:RP.057 complex conformation.

MD simulations were performed in the NVT ensemble at 37 °C and pH 7 using the CALVADOS suite.^37^ Simulations used a Langevin integrator with a time step of 10 fs and a friction coefficient of 0.01ps^-1^. Salt-screened electrostatic interactions were modelled using a dielectric constant of 74.2 and an ionic strength of 0.15 M. Simulations were conducted in a periodic box with dimensions of 40x40x40 nm. Simulations of RIPR, and RIPR with a single Fab, were run for a total of 5 μs. Simulations with two Fabs or larger complexes were run for 7.5 μs. Simulation convergence was measured by evaluating the mean distance between the RIPR-Body and the CTD centres of mass over the simulation time, as well as the similarity of the conformational space explored by the RIPR-Tail at different simulation lengths (**Figure S6**).

The coarse-grained bead model was implemented using parameters from the CALVADOS 3 multi-domain protein (MDP) release**^37^**, following the functional forms described in the original CALVADOS 3 publication. In all simulations, the RIPR-Body, the RIPR^CTD^ domain, the simulated Fabs, and the EGF domains in the RIPR-Tail, as defined using their UniProt definition, were treated as folded domains. Residues in these folded regions were restrained using a harmonic potential with a force constant of 700 N/m. In contrast, residue in the RIPR-Tail located between two consecutive EGF domains were considered disordered linkers. To capture the effect of Fabs on the flexibility of the RIPR**-**Tail, a scaled Lennard-Jones potential with a scaling coefficient of 30 kJ/mol was applied to selected residue pairs between the RIPR epitope and the Fab paratope. These pairs were identified in the cryo-EM structure by evaluating side-chain contacts, with a selection threshold of 4.5 Å between the centres of mass of the side chains being used (with the Cα atom being used for glycine).

The MDAnalysis suite**^60^** was used to process the MD trajectories and evaluate the different observables used during the study. These included the distance between the RIPR-Body and RIPR^CTD^ domain centres of mass, contact maps and the coordinates of the EGF domain centres of mass for the RIPR-Tail conformation analysis. For the latter, we extracted the distance matrix between the EGF centres of mass and used t-SNE^61^ to extract the embedding in the reduced space. Clustering of the structures was done using the Hierarchical Density-Based Spatial Clustering of Applications with Noise (HDBSCAN).^62^

### Quantification and statistical analysis

Data were analysed using GraphPad Prism version 10.4.1 for Windows (GraphPad Software Inc.).

